# Arabidopsis *At*GELP53 modulates polysaccharide acetylation and defense response through oligosaccharide-mediated signaling

**DOI:** 10.1101/2024.09.16.613205

**Authors:** Lavi Rastogi, Muhammad-Moazzam Hussain, Shivangi Tyagi, Kristi Kabyashree, Naman Kharbanda, Tushar Kanti Maiti, Vengadesan Krishnan, Gopaljee Jha, Aline Voxeur, Prashant Anupama-Mohan Pawar

## Abstract

*O*-acetylation is a crucial substitution found in hemicelluloses and pectin which are necessary for maintaining the flexibility and structural integrity of the cell. A balanced polysaccharide *O*-acetylation level is maintained by cell wall acetyl transferases and esterases present in different cell organelles. Specifically, the role of esterases in cell wall acetylation metabolism is less explored. Therefore, we investigated the role of *At*GELP53 from the GELP family of esterases or lipases. Here, we show that *At*GELP53 is localized in the plasma membrane. Analysis of *At*GELP53 overexpressing independent transgenic lines revealed a decrease in xyloglucan acetylation, changes in other polysaccharide acetylation and alteration in cell wall composition. The series of elicitor-based, transcriptomic and proteomic analyses in *At*GELP53 overexpressing lines suggested that oligosaccharide-mediated signalling activates the cell wall and defence-related genes primarily because of xyloglucan deacetylation. Furthermore, *At*GELP53 overexpression plants showed resistance against *Pseudomonas syringae* and *Ralstonia solanacearum* through activation of elicitor-mediated defense response. Overall, our findings outline the role of *At*GELP53 in polysaccharide acetylation, cell wall remodelling and defense through the activation of plant cell wall integrity.

## INTRODUCTION

Plant cell is surrounded by a dynamic cell wall covering which plays vital role in growth, development and immunity (De Lorenzo et al., 2019; Delmer et al., 2024). The plant cell wall is structurally built of a complex mixture of cellulose, hemicelluloses, pectin polysaccharides, and lignin polymer. The individual polysaccharide properties, cross-linking and interactions between components are governed by side chain substitutions (acetyl, methyl, glucuronic acid), present mainly on hemicelluloses and pectin, make the cell wall structure more complex and dynamic (Terrett & Dupree, 2019). These decorations, especially polysaccharide *O*-acetylation, the prevalent one, must be balanced for proper plant growth and development (Pawar et al., 2013; Qaseem & Wu, 2020; Zhong et al., 2024). Hemicelluloses such as xylan and xyloglucan are substituted with acetyl groups at varying positions in the backbone and side chain sugar moieties. The xylose residues in the xylan backbone are acetylated at *O*-2 and/or *O*-3 positions and side chain galactose residues in xyloglucan are mainly acetylated at *O*-6, *O*-5 and *O*-4 positions. Matrix polysaccharide pectin includes homogalacturonan (HG), rhamnogalacturonan-I (RG-I), and rhamnogalacturonan-II (RG-II) which are acetylated at *O*-2 and/or *O*-3 positions of backbone galacturonic acid residues in HG and RG-I whereas *O*-acetylation was found at 2-*O*-methyl-fucose in side chain residues in RG-II (Shahin et al., 2023). These polysaccharides are synthesized in the Golgi and substituted with acetyl groups via various proteins localized in the Golgi membrane and further modifications may be carried out in the cell wall (Pauly & Ramirez, 2018; Rastogi et al., 2022; Zhong et al., 2020). However, the post-synthetic modification in polysaccharide *O*-acetylation is relatively less explored. Also, factors or enzymes regulating acetylation homeostasis and their role post-synthesis modification are not studied.

Previous findings suggested that hypo-and hyper-acetylation of cell wall polysaccharides can disrupt plant growth (Manabe et al., 2013; Schultink et al., 2015; Yuan et al., 2013; Zhang et al., 2017; Zhang et al., 2019). The group of protein families including REDUCED WALL ACETYLATION (RWA), ALTERED XYLOGLUCAN 9 (AXY9), and TRICHOME BIREFRINGENCE-LIKE (TBL) family are involved in the decoration of acetyl groups on polysaccharide chain. Arabidopsis quadruple *rwa1/2/3/4*, *axy9*, and *tbl29* mutants show a reduction in polysaccharide acetylation by about 60%, 80%, and 60%, respectively, with severe growth defects (Manabe et al., 2013; Schultink et al., 2015; Yuan et al., 2013). On the other hand, the cases related to hyper-acetylation were observed in rice mutants with defects in the GDSL esterase/lipase family members that function antagonistically to acetyltransferases. Mutation in Golgi localized *BRITTLE LEAF SHEATH 1* (*BS1*) and *DEACETYLASE ON ARABINOSYL SIDECHAIN OF XYLAN 1* (*DARX1*) GDSL esterases led to elevated levels of polysaccharide acetylation around 133% and 14% respectively and both rice mutants (*bs1 and darx1*) are dwarf (Zhang et al., 2017; Zhang et al., 2019). Hence, a balanced polysaccharide *O*-acetylation level is required to maintain overall cell wall architecture which is governed by both polysaccharide transferases and esterases.

Post-synthesis modification is tolerated by plants as either overexpression of *An*AXE1 or *At*AXE1 reduces xylan acetylation without affecting plant growth. Other compensatory changes in gene expression of acetylation pathway genes and an increase in acetyl content of another polysaccharide were observed (Pawar et al., 2016; Rastogi et al., 2022). This suggests that plants can compensate for the post-synthesis acetylation loss in cell wall polysaccharides and maintain the required amount of polysaccharide *O*-acetylation essential for plant growth. Interestingly, this raises an important question: how do plants sense acetylation changes in the cell wall? The simplest hypothesis could be that due to the loss of acetyl groups by the action of acetyl esterases, there might be a change in the apoplastic pH through which plants sense acetylation changes in the cell wall. Another explanation could be that because of the loss of acetyl groups, polysaccharides become prone to the action of cell wall endohydrolases and release oligosaccharides, which might act as signaling molecules or elicitors that trigger a signaling cascade through binding to some specific cell wall receptors. This further raises another exciting question, i.e., how do plants compensate for the acetylation loss? For compensation, plants might alter polysaccharide acetylation by altering the expression of cell wall acetylation genes, and this is how they might maintain acetylation homeostasis in the cell wall. In this manuscript, we tried to address these questions by studying a member of the GELP family. In Arabidopsis, many GELPs are characterized having role in lipid metabolism, but very few of them are characterised for their role in cell wall metabolism. So far, a FUCOSIDASE (*At*FXG1/*At*GELP33), CUTICLE DESTRUCTING FACTOR 1 (CDEF1/*At*GELP83), and ACETYL XYLAN ESTERASE (*At*AXE1/*At*GELP7) are characterized that are known to have functions in the plant cell wall modification (De La Torre et al., 2002; Rastogi et al., 2022; Takahashi et al., 2010). Therefore, in this study, we are characterizing *At*GELP53 from clade Id of Arabidopsis GELP family by overexpressing it in Arabidopsis and studying changes in cell wall modifications, especially acetylation, to further understand the maintenance in cell wall integrity in this model plant.

In this study, we found that *A*tGELP53 is a localized in plasma membrane and its overexpression in Arabidopsis reduces xyloglucan acetylation which was further validated by docking studies. In contrast, pectin and xylan polysaccharides in *A*tGELP53 overexpressor lines showed more acetylation, suggesting a compensatory effect of reduced acetylation in xyloglucan. Elicitor studies and proteomics analyses revealed that elicitor-mediated changes in ROS production and induction in plant defence marker, cell wall integrity and acetylation pathway genes. All these changes might have led to resistance against *Pseudomonas syringae* and *Ralstonia solanacearum* in *A*tGELP53 overexpressing lines as compared to wild-type plants.

## RESULTS

### Characterized proteins in the Arabidopsis GELP family

The GELP family, known as the GDSL esterase/lipase protein family, belongs to the SGNH hydrolase superfamily with Serine (S), Glycine (G), Asparagine (N), and Histidine (H) amino acids conserved in four blocks I, II, III, and V respectively, in all GELP amino acid sequences. These GELP members can perform multiple functions in plants because they have a very flexible enzyme active site that provides specificity to a variety of substrates. The GDS motif at the N-terminal and DXXH motif at the C-terminal are also conserved in this protein family which are important for their catalytic activity (Akoh et al., 2004). In *Arabidopsis thaliana*, the GELP family has not been explored much in terms of its role in cell wall metabolism instead, some of these proteins have been characterized as having role in lipid metabolism (Agee et al., 2010; Jancowski et al., 2014; Lee et al., 2009; Mayfield et al., 2001; Nagano et al., 2008; Oh et al., 2005). Phylogenetic tree analysis revealed that Arabidopsis GELP family with 105 members were categorized into two clades (Figure 1A). The clade I is further sub-categorized into ten subclades, i.e., clade Ia, Ib, Ic, Id, Ie, If, Ig, Ih, Ii, and Ij. The characterized GELP proteins are highlighted in each clade. We also made a list of characterized GELP proteins with their probable function (Figure 1B). In our previous finding, we characterised *AtGELP7* from clade Id as acetyl xylan esterase (*AtAXE1*), suggesting clade Id members are involved in polysaccharide de-esterification (Rastogi et al., 2022). Therefore, in the present study, we performed functional characterization of *At*GELP53 and elucidated its role in cell wall metabolism.

**Figure 1.**
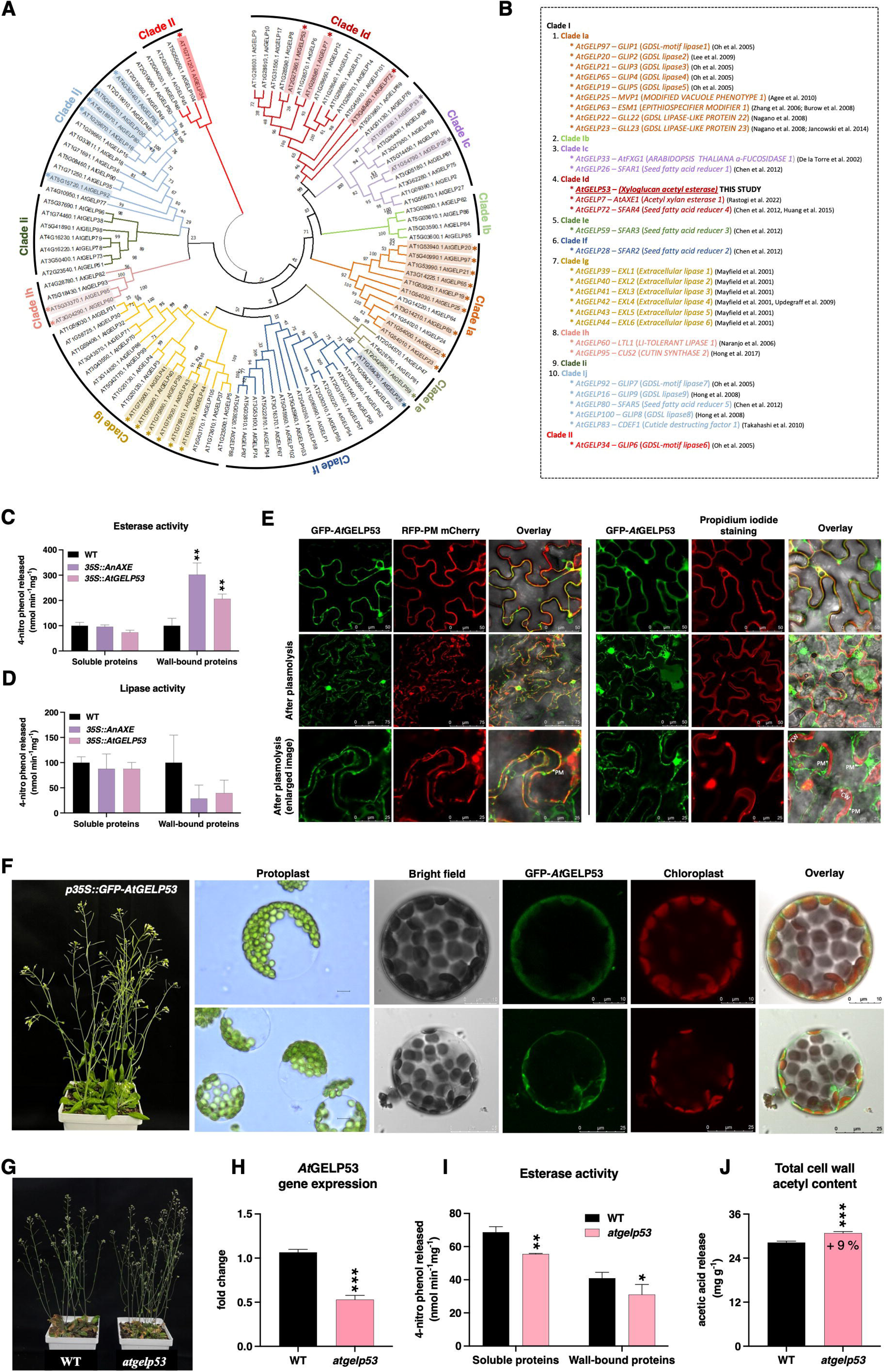
*At*GELP53 from the Arabidopsis GELP family is a plasma membrane-localized acetyl esterase. **(A)** Phylogenetic tree of Arabidopsis GELP family generated by MEGA11 software using the maximum likelihood (ML) method with 1000 bootstraps. The tree included 105 GELP proteins segregated into clades based on their sequence similarity. **(B)** A List displaying annotated GELP proteins from each clade and *At*GELP53 is highlighted in clade Id. **(C-D)** Agrobacterium carrying *35S::AtGELP53* transiently expressed in Nicotiana leaves by infiltration and esterase **(C)** and lipase **(D)** activity was estimated by incubating soluble and wall-bound proteins with 4-nitrophenyl acetate and 4-nitrophenyl palmitate substrates, respectively. The released 4-nitro phenol was detected at 400 nm by spectrophotometer and represented on Y-axis. Data represents mean ± SE, *n* = 3-4 biological replicates, Student’s t-test at \*\*\*\**p* ≤ 0.001, \*\*\**p* ≤ 0.01, \*\**p* ≤ 0.05, * *p* ≤ 0.1. **(E)** *At*GELP53 was tagged with N-terminal GFP and co-infiltrated with plasma membrane RFP marker in Nicotiana epidermal leaf cells and after 2^nd^-day post infiltration, the leaf sections were visualized under confocal microscope. The leaf sections were also stained with a cell wall stain i.e., propidium iodide, and pictures were captured before and after plasmolysis in 30% glycerol with and without cell wall stain. PM; plasma membrane, CW; cell wall. **(F)** *At*GELP53 GFP stable Arabidopsis plants were generated as shown in the picture and protoplasts were isolated from their leaves and directly imaged under light and confocal microscope. Chloroplast autofluorescence was visualized in the red channel. **(G)** Picture of 6 weeks-old wild-type Arabidopsis plant with *atgelp53* homozygous mutant plant. **(H)** RNA level expression of *At*GELP53 gene in *atgelp53* homozygous mutant plant by qRT-PCR. *ACTIN* was used as reference gene. Data represents mean ± SE, *n* = 3-4 biological replicates, Student’s t-test at \*\*\*\**p* ≤ 0.001, \*\*\**p* ≤ 0.01, \*\**p* ≤ 0.05, * *p* ≤ 0.1. **(I)** Esterase activity was detected in soluble and wall-bound protein fractions with 4-nitrophenyl acetate as substrate in wild-type and *atgelp53* homozygous mutant. The released 4-nitro phenol was detected at 400 nm by spectrophotometer and represented on Y-axis. Data represents mean ± SE, *n* = 3-4 biological replicates, Student’s t-test at \*\*\*\**p* ≤ 0.001, \*\*\**p* ≤ 0.01, \*\**p* ≤ 0.05, * *p* ≤ 0.1. **(J)** Alcohol insoluble residues (AIR) were extracted from *atgelp53* homozygous mutant plants, de-esterified using 1M NaOH and the total cell wall acetyl content was measured by acetic acid kit. Data represents mean ± SE, *n* = 3-4 biological replicates, Student’s t-test at \*\*\*\**p* ≤ 0.001, \*\*\**p* ≤ 0.01, \*\**p* ≤ 0.05, * *p* ≤ 0.1.

### *At*GELP53 is localized in the plasma membrane and its mutant shows reduced esterase activity

*AtGELP53* was cloned into a vector containing CAMV 35S constitutive promoter and expressed transiently in *Nicotiana benthamiana* leaves. Both esterase and lipase activities were tested using synthetic substrates by isolating protein from infiltrated leaves and *Aspergillus niger* acetyl xylan esterase (*AnAXE1*) was used as a control for esterase activity (Pawar et al., 2016). *At*GELP53 showed increased esterase activity in the wall-bound protein fraction similar to *An*AXE1 positive control but did not show increased lipase activity (Figure 1C and 1D). To check subcellular localization, GFP-tagged *At*GELP53 was co-infiltrated with an RFP-tagged plasma membrane protein (PIP 2A) and an overlay image showed co-localization of *At*GELP53 with the plasma membrane marker (Figure 1E). Propidium iodide was used to stain the cell wall and *At*GELP53 GFP signal was also found overlaying with propidium iodide suggesting its localization either in the plasma membrane or in the cell wall. Additionally, the plasmolysis experiment confirmed *At*GELP53 localization in the plasma membrane. To confirm this further, Arabidopsis stable GFP lines of *At*GELP53 were generated, the protoplasts were isolated and directly visualized under the confocal microscope (Figure 1F). The GFP signal was clearly visible in the plasma membrane. The transient and stabled expression experiments demonstrated that *At*GELP53 is a plasma-membrane localized esterase.

To further determine the function of *At*GELP53, we isolated and characterized its homozygous T-DNA mutant named *atgelp53* (Figure S1A and S1B). Morphologically, the *atglep53* looked similar to the wild-type plant (Figure 1G). The expression of *AtGELP53* at the RNA level was found to be 50 % reduced in the stem of the *atglep53*, suggesting that it is a knockdown mutant (Figure 1H). The esterase activity was also reduced in both soluble and wall-bound protein fractions in *atglep53* (Figure 1I). The total cell wall acetyl content in *atgelp53* mutant was increased by 9% compared to the wild-type (Figure 1J). To further investigate acetylation changes in *atgelp53*, we digested AIR with xyloglucanase, pectinase, and xylanase separately and released oligosaccharides were analyzed by liquid chromatography-mass spectrometry (LC-MS). In enzymatic oligosaccharide profiling data, we observed slight changes in the pattern of pectin and xylan oligosaccharide release in the *atgelp53* mutant, but the xyloglucan oligosaccharide profile was similar in mutant and wild-type (Figure S2A-C).

### *At*GELP53 is expressed mainly in primary cell wall forming tissue and *35S::AtGELP53* **overexpression lines showed a decrease in total cell wall acetyl content**

To check the expression of *At*GELP53 in different plant tissues, we generated Arabidopsis stable line containing *AtGELP53* promoter fused with GLUCURONIDASE (GUS). *At*GELP53 expression was primarily found in rosette leaves, inflorescence, and roots, suggesting it is active mainly in primary tissue (Figure 2A). To explore changes in cell wall polysaccharide acetylation and to further characterize *At*GELP53, three independent transgenic lines under 35S constitutive promoter were generated, i.e., *At*GELP53-3, *At*GELP53-4, and *At*GELP53-5 (Figure 2B). Morphologically, *At*GELP53 transgenic plants showed more height compared to the wild-type plant (Figure 2C). All three lines showed higher expression of GELP53 as compared to wild-type in both leaf and stem tissue (Figure 2D and 2E). The esterase activity was significantly higher in soluble and wall-bound protein fractions in both leaf and stem tissue in *At*GELP53 overexpressing (OE) lines (Figure 2F and 2G). As the GUS stable line also showed expression of *At*GELP53 in roots; therefore, root morphology was also studied (Figure 2H). The root length of 12-day-old seedlings of transgenic lines was more as compared to wild-type plants (Figure 2I). Subsequently, all these tissues (leaf, stem, root and seedlings) showed a significant reduction in acetyl content in *At*GELP53 OE lines with an average of 7.3% in leaf, 11.5% in stem, 12% in roots, and 20% in seedlings (Figure 2J-L and Figure S3). This data suggested that *At*GELP53 is a polysaccharide acetyl esterase and may be prominently functioning in primary cell wall-forming tissue under normal conditions.

**Figure 2.**
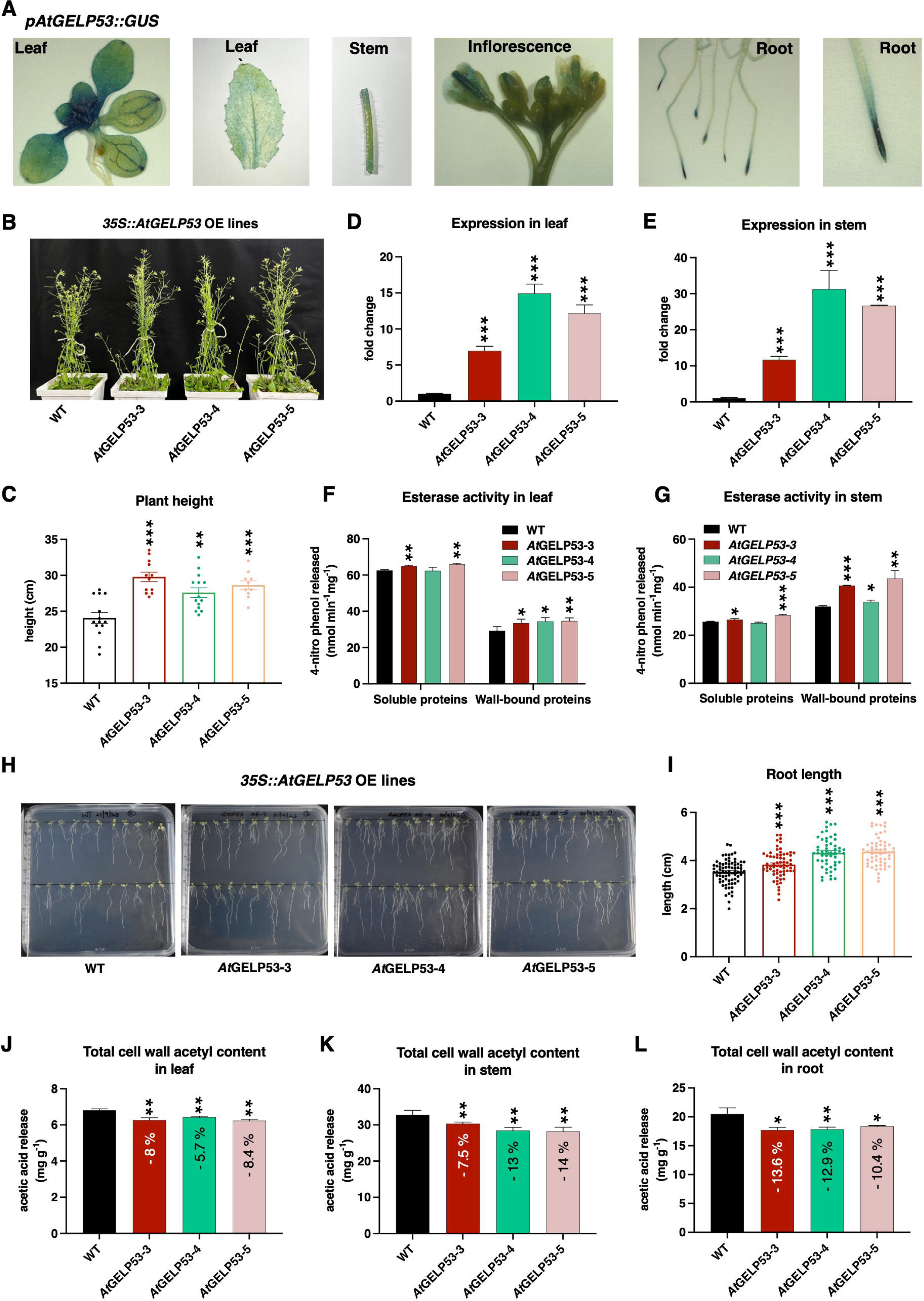
Overexpression of *At*GELP53 improves growth characteristics and reduces total cell wall acetylation. **(A)** The gene expression of *AtGELP53* was detected in different plant tissues such as leaf, stem, roots, and inflorescence of *pAtGELP53::GUS* stable Arabidopsis line by promoter-GUS activity using X-Gluc as substrate. The pictures were captured under the light microscope. **(B)** The picture of 5-6-week-old wild-type Arabidopsis plant with three independent transgenic lines i.e., *At*GELP53-3, *At*GELP53-4, and *At*GELP53-5 overexpressing *AtGELP53* under 35S constitutive promoter. **(C)** The height of *35S::AtGELP53* overexpression plants was measured and compared to the height of wild-type Arabidopsis plants. Data represents mean ± SE, *n* = 10-15 biological replicates, Student’s t-test at \*\*\*\**p* ≤ 0.001, \*\*\**p* ≤ 0.01, \*\**p* ≤ 0.05, * *p* ≤ 0.1. **(D-E)** The expression of *AtGELP53* gene was checked in leaf **(D)** and stem **(E)** of *35S::AtGELP53* OE lines by qRT-PCR. *ACTIN* was used as reference gene. Data represents mean ± SE, *n* = 3-4 biological replicates, Student’s t-test at \*\*\*\**p* ≤ 0.001, \*\*\**p* ≤ 0.01, \*\**p* ≤ 0.05, * *p* ≤ 0.1. **(F-G)** Esterase activity was detected in soluble and wall-bound protein fractions with 4-nitrophenyl acetate as substrate in the leaf **(F)** and stem **(G)** of *35S::AtGELP53* OE lines. The released 4-nitro phenol was detected at 400 nm by spectrophotometer and represented on Y-axis. Data represents mean ± SE, *n* = 3-4 biological replicates, Student’s t-test at \*\*\*\**p* ≤ 0.001, \*\*\**p* ≤ 0.01, \*\**p* ≤ 0.05, * *p* ≤ 0.1. **(H)** The picture shows 12-day-old seedlings of *35S::AtGELP53* OE lines grown on Murashige and Skoog (MS) media. **(I)** The root length of *35S::AtGELP53* OE seedlings was measured and compared to the root length of wild-type seedlings. Data represents mean ± SE, *n* = 20-22 biological replicates, Student’s t-test at \*\*\*\**p* ≤ 0.001, \*\*\**p* ≤ 0.01, \*\**p* ≤ 0.05, * *p* ≤ 0.1. **(J-L)** Alcohol insoluble residues (AIR) were extracted and de-esterified using 1M NaOH and the acetyl content was measured in leaf **(J)**, stem **(K)**, and roots **(L)** of *35S::AtGELP53* OE lines in comparison with wild-type plant by acetic acid kit. Data represents mean ± SE, *n* = 3-4 biological replicates, Student’s t-test at \*\*\*\**p* ≤ 0.001, \*\*\**p* ≤ 0.01, \*\**p* ≤ 0.05, * *p* ≤ 0.1.

### AtGELP53 could be involved in xyloglucan deacetylation

To elucidate the primary function of *At*GELP53 in polysaccharide de-esterification, the polysaccharides from leaf AIR was sequentially extracted according to the flow chart (Figure S4) and acetyl content was measured in each fraction. The pectin-rich fraction I (Ammonium formate soluble fraction) and fraction II (non-methylesterified homogalacturonans) showed increased acetyl content in *At*GELP53 OE lines compared to wild-type plant (Figure 3A and 3B). The depectinized pellet was digested with xyloglucanase and supernatant containing xyloglucan-rich fraction showed drastically reduced acetyl content with an average of 44% in *AtGELP53* OE lines (Figure 3C). The remaining pellet showed increased acetyl content in *AtGELP53* OE lines (Figure 3D). Altogether this data shows that *At*GELP53 overexpression reduces acetylation only on xyloglucan polysaccharide or co-extracted polysaccharides and not on low methylesterified pectin or cellulose enriched-fractions which suggests that *At*GELP53 could be specifically deacetylating xyloglucan and acting as xyloglucan acetyl esterase. On the contrary, increased acetylation in other fractions in *At*GELP53 OE lines could be a compensatory effect.

**Figure 3.**
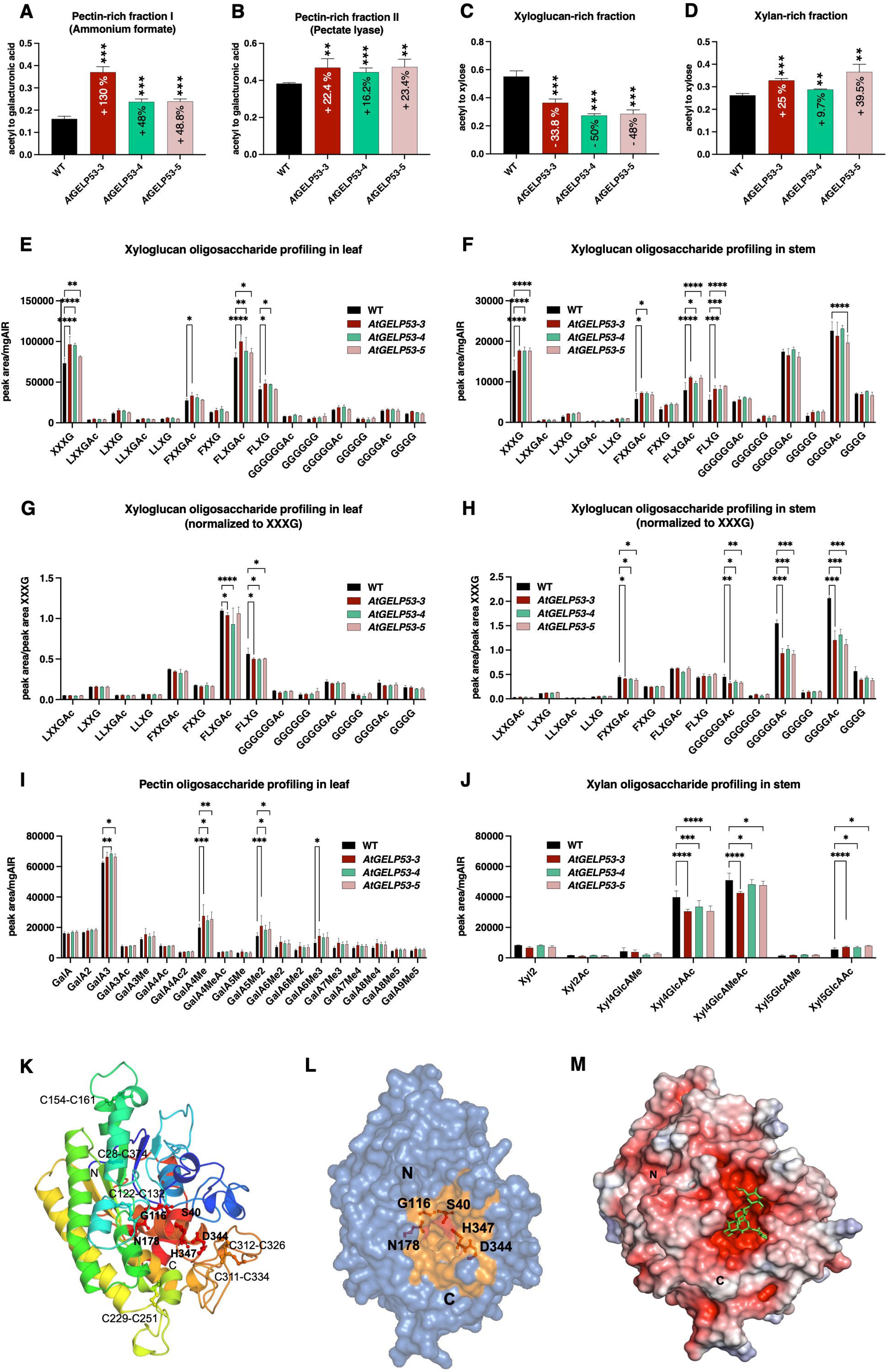
*At*GELP53 is involved in the deacetylation of xyloglucan. **(A-D)** Different cell wall polysaccharides were sequentially extracted from leaf AIR of wild-type and *35S::AtGELP53* OE lines and acetyl content was analyzed in pectin rich ammonium formate fraction I **(A)**, pectin rich pectate lyase fraction II **(B)**, xyloglucan rich fraction **(C)**, and xylan rich pellet fraction **(D)** and normalized with their respective sugar content. Data represents mean ± SE, *n* = 3-4 biological replicates, Student’s t-test at \*\*\*\**p* ≤ 0.001, \*\*\**p* ≤ 0.01, \*\**p* ≤ 0.05, * *p* ≤ 0.1. **(E-J)** Leaf and stem AIR of wild type and *35S::AtGELP53* OE lines were digested with xyloglucanase and released oligosaccharides were detected by liquid chromatography-mass spectrometry (LC-MS) in leaf **(E)** and stem **(F)** and peak area was also normalized to peak area of XXXG in leaf **(G)**, and stem **(H)**. AIR was also digested with polygalacturonase (leaf) **(I)**, and xylanase (stem) **(J)**, and released oligosaccharides were detected by LC-MS. X, Xyl-xylose, G-glucose, L-galactose, F-fucose, GalA-galacturonic acid, GlcA-glucuronic acid, Ac-*O*-acetyl, Me-*O*-methyl. Data represents mean ± SE, *n* = 3-4 biological replicates, Student’s t-test at \*\*\*\**p* ≤ 0.001, \*\*\**p* ≤ 0.01, \*\**p* ≤ 0.05, * *p* ≤ 0.1. **(K)** A Ribbon diagram of the overall architecture of *At*GELP53 predicted by AlphaFold2 server. The structure is shown in a rainbow color style, and the N- and C-terminals are labelled. Active site and oxyanion hole residues are indicated at the active site cleft. The disulfide bridges such as Cys28-Cys374, Cys122-Cys132, Cys154-Cys161, Cys229-Cys251, Cys311-Cys334, and Cys312-Cys326, are also labelled. **(L)** The surface representation of the *At*GELP53 protein shows an active site cleft with the catalytic triad having Ser40, Asp344, and His347 (red sticks). Residues involved Gly116 and Asn178 involved in oxyanion hole formation with Ser40 also shown (red sticks). **(M)** Electrostatic surface representation of *At*GELP53 showing large size negatively charged active site cleft with a docked xyloglucan moiety (green sticks). The docking was done through AutoDock Vina software.

To further confirm that in *At*GELP53 OE lines xyloglucans were less acetylated, we performed an enzymatic fingerprinting approach. Alcohol insoluble residue (AIR) of *At*GELP53 was subjected to digestion with xyloglucanase, and released oligosaccharides were detected by LC-MS (Figure 3E and 3F). In xyloglucan oligosaccharide profiling, we first observed that more oligosaccharides, and particularly XXXG and fucosylated oligosaccharides, were released from leaf and stem AIRs of *At*GELP53 OE lines suggesting that *At*GELP53 OE lines were enriched in xyloglucans compared to the wild type. Interestingly, we also observed that more cello-oligosaccharides were released from stems compared to leaves and that their acetylated equivalent were also detected. When the peak areas of the more substituted xyloglucan fragments and the cello-oligosaccharides were normalized to the peak area of XXXG, we observed significantly less acetylated and non-acetylated FLXG in the leaves of *At*GELP53 OE lines, as well as a tendency towards fewer acetylated cello-oligosaccharides (Figure 3G). In the stems, we observed a trend towards fewer acetylated FXXG and significantly fewer acetylated cello-oligosaccharides in *At*GELP53 OE lines compared to the wild-type (Figure 3H). Together, the observations of reduced acetylation in xyloglucan and cello-oligosaccharides, when normalized to XXXG content, are consistent with the previously observed lower acetyl content in xyloglucan-rich fraction relative to xylose.

Furthermore, the fine structures of homogalacturonan and xylan were also assessed using enzymatic fingerprinting. Leaves and stems from *At*GELP53 OE lines were subjected to digestion with polygalacturonase, which digests low and moderately methylesterified pectins (unlike the previously used pectate lyase), and xylanase, respectively (Figure 3I and 3J). In leaves, more pectic oligosaccharides were released from *At*GELP53 OE lines, suggesting that either the homogalacturonans were more digestible (i.e., more demethylesterified) or that *At*GELP53 cell walls are enriched in pectins compared to the wild type. However, no differences in acetylated oligosaccharides were observed, suggesting that the increased pectic acetylation affects the highly soluble and non-methylesterified pectic populations. Next, stem AIR were subjected to xylanase, an enzyme whose activity is known to be partially inhibited by xylan acetylation (Biely et al., 1986). In the xylan oligosaccharide profiling, we observed that fewer oligosaccharides were released from the *At*GELP53 OE lines, which was expected given the higher acetyl content relative to xylose in the pellet. Additionally, while fewer glucuronosylated tetraxylosyl oligosaccharides were detected, more pentaxylosyl glucuronosylated oligosaccharides were observed, suggesting that the modification of the xylan decoration pattern might be a secondary effect of *At*GELP53 activity. In detail, shorter oligosaccharides, i.e., Xyl2 and Xyl2Ac levels were not changed but shorter-acetylated oligosaccharide, i.e., Xyl4GlcAAc level was less, whereas longer-acetylated and methylated oligosaccharides, i.e., Xyl5GlcAAc, and Xyl5GlcAMe levels were more in transgenic lines. Based on the pattern of oligosaccharides released after enzymatic digestion, these results demonstrated that xyloglucanase accessibility towards xyloglucan is more because of less xyloglucan acetylation and as a result, there is more release of shorter oligosaccharide in *At*GELP53 OE lines. Whereas polygalacturonase and xylanase accessibility towards pectin and xylan, respectively, are less because of more pectin and xylan acetylation and as a result, there is less release of shorter oligosaccharide in *At*GELP53 OE lines.

Overall, reduced acetylation in xyloglucan-rich fraction and altered xyloglucan oligosaccharide profile in GELP53 OE lines indicated *At*GELP53 as xyloglucan acetyl esterase.

### Structural analysis of *At*GELP53 and substrate binding

The full-length *At*GELP53 comprises 394 amino acids, including a signal peptide at the N-terminal (Figure S5A). The GDS and DXXH motifs are located near the N- and C-terminals within a single catalytic domain. Structurally, *At*GELP53 features a central β-sheet of five β-strands (β2-β1-β3-β4-β5) flanked by α-helices, forming the α/β topological fold typical of the GDSL esterase/lipase hydrolase superfamily (Figure S5B) (Akoh et al., 2004). The *At*GELP53 structure resembles the SGNH-hydrolase family, which shares five consensus blocks (Lo et al., 2003). Like GDSL enzymes, *At*GELP53 contains a flexible active site to recognize various substrates, with the catalytic triad comprising Ser40, Asp344, and His347. A DALI search for structural homologs of *At*GELP53 identified many hydrolase structures with significant Z-scores but limited sequence identity (<25%), confirming that *At*GELP53 belongs within the SGNH-hydrolase family (Figure S5C) (Table S1) (Holm & Rosenstrom, 2010). Compared to other known structures, *At*GELP53 has more disulfide bridges, many located on loops connecting secondary structural elements (Figure 3K). Notably, the disulfide bridge loop Cys311-Cys334 likely regulates substrate access to the active site, akin to the disulfide loop in lysophospholipase A, processed by zinc metalloproteinase ProA to enhance activity (Hiller et al., 2024). In *At*GELP53, the Ser40 residue (nucleophile) is part of the block I GDSI motif near the N-terminal, while block II includes Gly116, block III features Asn178, and block V contains Asp344 and His347, essential for forming the catalytic triad and oxyanion hole (Figure 3K and 3L). The active site cleft, located at the top of the central β-sheet, is surrounded by many hydrophilic and hydrophobic residues (Ile41, His347, Asp344, Ile346, Tyr287, Asp39, Glu174, Asn224, Asn178, Gly177, Asn181, Leu227, His262). The oxyanion hole, crucial for stabilizing the oxyanion generated during substrate cleavage is formed by Ser40, Gly116, and Asn178 (Akoh et al., 2004).

Molecular docking with xyloglucan, a possible substrate identified from *At*GELP53 overexpression studies, revealed that *At*GELP53 can accommodate this substrate at its large acidic active site cleft through various interactions (Figure 3M and Figure S5D and E). Asn153 and Pro201 participate in hydrogen bonding with xyloglucan, while hydrophobic interactions involve residues such as Ile41, Phe76, Ala117, Phe185, Phe225, Leu227, Cys229, Leu291, Ile346, Met348, and Tyr353. Polar (Ser40, Asn181, Ser230, Asn342, and His347), charged residues (Asp39, Glu58, and Glu174), and glycine residues (Gly115, Gly116, and Gly177), also participate in substrate binding. Despite having limited sequence identity with known structures, the *At*GELP53 resembles the SGNH-hydrolase family with a conserved catalytic triad and a large-size flexible active site that can accommodate xyloglucan.

### Polysaccharide acetylation level in *At*GELP53 overexpression plants negatively affects polysaccharide extractability and digestibility

To further understand the effect of changes in acetylation levels on xyloglucan, pectin, and xylan in *At*GELP53 OE lines, we immunolabeled cell wall polysaccharides with glycan-specific antibodies. The stem cross sections of wild-type and *At*GELP53 OE lines were stained with toluidine blue, and images were used as a representative to show the whole stem anatomy (Figure 4A). These sections were immunolabeled with LM15, LM20, LM10, and LM11 antibodies specific for different wall polysaccharides (Figure 4A). The LM15 antibody recognizes specifically the XXXG motif in xyloglucan, and LM15 signals were detected more in *At*GELP53 OE lines. This result correlated with xyloglucan oligosaccharide profiling data, which showed more release of shorter XXXG oligosaccharides. LM20 antibody recognizes methyl-esterified homogalacturonan, i.e., pectin epitopes, and the signals were detected more in *At*GELP53 OE lines correlating with the detection of more methyl-esterified fragments in pectin oligosaccharide profiling. The epitopes of LM10 and LM11 antibodies are unsubstituted xylan and substituted xylan, respectively. LM10 signals were detected less in *At*GELP53 transgenic lines, whereas LM11 signals were detected more compared to wild-type, suggesting the presence of more substituted xylan in *At*GELP53 OE lines.

**Figure 4.**
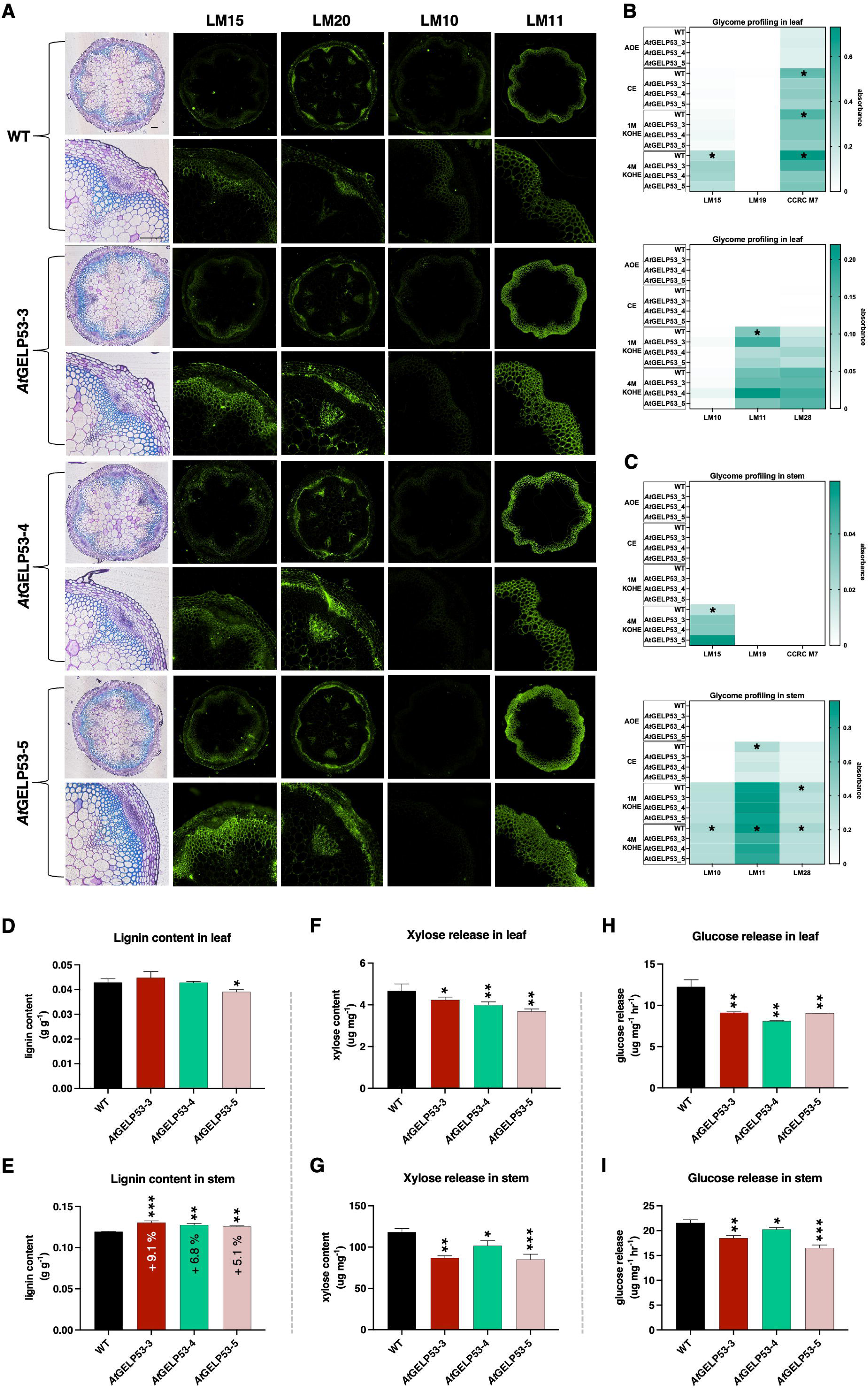
Cell wall glycan immunolocalization, extractability and digestibility are negatively correlated with polysaccharide acetylation profile in *At*GELP53 overexpression lines. **(A)** Transverse sections of main stems of wild-type and *35S::AtGELP53* OE lines stained with toluidine blue and immunolabelled with LM15, LM20, LM10 and LM11 antibodies that targets XXXG motif in xyloglucan, methyl-esterified pectin, unsubstituted xylan and substituted xylan respectively. The images were captured under Nikon microscope. Scale bar: 100 µm. **(B-C)** Alcohol insoluble residues (AIR) from leaf **(B)** and stem **(C)** of wild-type plants and *35S::AtGELP53* OE lines were sequentially treated with ammonium oxalate, carbonate, 1M potassium hydroxide, 4M potassium hydroxide buffers and different extracts were collected as ammonium oxalate extract (AOE), carbonate extract (CE), 1M potassium hydroxide extract (1MKOHE), and 4M potassium hydroxide extract (4MKOHE) and immunolabelled with LM15, LM19, CCRC M7, LM10, LM11, and LM28 antibodies in ELISA plate. The chemiluminescence was detected at 450 nm with a background reading at 655 nm in spectrophotometer. Data represents mean ± SE, *n* = 3-4 biological replicates, Student’s t-test at \*\*\*\**p* ≤ 0.001, \*\*\**p* ≤ 0.01, \*\**p* ≤ 0.05, * *p* ≤ 0.1. **(D-E)** Lignin content was measured by acetyl bromide soluble lignin (ABSL) method in leaf **(D)** and stem **(E)** AIR of wild-type plants and *35S::AtGELP53* OE lines. Data represents mean ± SE, *n* = 3-4 biological replicates, Student’s t-test at \*\*\*\**p* ≤ 0.001, \*\*\**p* ≤ 0.01, \*\**p* ≤ 0.05, * *p* ≤ 0.1. **(F-G)** Enzymatic digestion of leaf **(F)** and stem **(G)** AIR with xylanase and xylose release was detected by xylose kit. Data represents mean ± SE, *n* = 3-4 biological replicates, Student’s t-test at \*\*\*\**p* ≤ 0.001, \*\*\**p* ≤ 0.01, \*\**p* ≤ 0.05, * *p* ≤ 0.1. **(H-I)** Saccharification efficiency was calculated by measuring glucose release in both leaf **(H)** and stem **(I)** AIR of wild type and *35S::AtGELP53* OE lines. Data represents mean ± SE, *n* = 3-4 biological replicates, Student’s t-test at \*\*\*\**p* ≤ 0.001, \*\*\**p* ≤ 0.01, \*\**p* ≤ 0.05, * *p* ≤ 0.1.

Alteration in acetyl modifications on individual polysaccharides in *At*GELP53 OE lines can be further understood through polysaccharide extractability assays such as ELISA-based glycome profiling. Initially, the cell wall fractionation was done by treating leaf and stem AIR of wild-type and transgenic plants with buffer and alkaline reagent and different extracts were isolated as ammonium oxalate extract (AOE), carbonate extract (CE), 1M potassium hydroxide extract (1MKOHE), and 4M potassium hydroxide extract (4MKOHE). These extracts were then subjected to ELISA plate to detect different polysaccharides using wall glycan-specific antibodies (Figure 4B and 4C). Xyloglucan polysaccharides were detected in 4MKOHE using a xyloglucan-specific antibody i.e., LM15, and showed more extractability in both leaf and stem tissue of *At*GELP53 OE lines compared to wild-type plants. Pectin polysaccharides were detected in CE, 1MKOHE, and 4MKOHE using a pectin-specific antibody i.e., CCRC M7, in leaf only and showed less extractability in transgenic plants. Xylan polysaccharides were detected in 1MKOHE and 4MKOHE using xylan-specific antibodies i.e., LM10, LM11, and LM28, in specifically stem and showed less extractability in *At*GELP53 OE lines compared to wild-type. These results revealed that xyloglucan extractability is higher, whereas pectin and xylan extractability is lower in transgenic lines as compared to wild-type.

To further analyze any changes in the cell wall composition that might have caused because of changes in the acetyl content, we measured the hemicellulosic sugar content by ion chromatography. We did not see much change in the individual hemicellulosic monosugars such as xylose, arabinose, rhamnose, fucose, galactose, mannose, and glucose content in leaf tissue but found slight decrease in xylose content and increase in other hemicellulosic monosugars except glucose in stem tissue of *At*GELP53 OE lines (Table S2). We also measured pectin, cellulose, and lignin content in the leaf and stem tissue of these lines and found that pectin and cellulose contents were similar in wild-type and transgenic lines, but there was a significant increase in lignin content in the stem of *At*GELP53 OE lines (Figure S6A-D and Figure 4D and 4E). Changes in the acetyl content and cell wall composition affects the digestibility of polysaccharide. So, we further tested xylan digestibility by digesting leaf and stem AIR with xylanase and measured the xylose release, which was found to be reduced in *At*GELP53 OE lines (Figure 4F and 4G). Xylan digestibility data correlated with xylan oligosaccharide profile and xylan acetyl content. Cellulose digestibility was also tested by using an enzyme cocktail having cellulase and β-glucosidase, and glucose release was measured, which was reduced in both the leaf and stem of transgenic lines compared to wild-type (Figure 4H and 4I). Overall, this data suggested that polysaccharide acetylation can negatively affect extractability and digestibility.

### Expression of several cell wall-associated genes was altered in *At*GELP53 OE lines

To correlate changes in the cell wall of *At*GELP53 OE lines, the expression of major cell wall-related genes involved in polysaccharide acetylation, lignin biosynthesis, cell wall integrity, and cell wall endohydrolases were evaluated. Initially, we evaluated the expression of key polysaccharide acetyltransferase genes belonging to TBL family, i.e., *TBL27*, *TBL10*, and *TBL29*, which are specifically known to acetylate xyloglucan, pectin, and xylan, respectively (Gille et al., 2011; Stranne et al., 2018; Yuan et al., 2013). The expression of *TBL27* was not significantly changed, but *TBL10* and *TBL29* expression were significantly upregulated in *At*GELP53 OE lines (Figure 5A-C). The upregulation of pectin acetyltransferase, i.e., *TBL10* and xylan acetyltransferase, i.e., *TBL29* correlating with the increased acetylation on pectin and xylan in transgenic lines. Apart from acetyltransferase genes, the expression of other genes involved in cell wall acetylation such as *RWAs*, *AXY9* and *ATP CITRATE LYASES* (*ACLA*) were also tested. In the leaf of *At*GELP53 OE lines, *RWA4* and *ACLA3* whereas in stem *RWA1*, *RWA3*, *ACLA1*, *ACLA2* and *AXY9* gene expressions were significantly upregulated (Figure 5D). As described above, lignin content was significantly higher in stem tissue of *At*GELP53 OE lines, so we checked the expression of lignin biosynthesis genes such as *PAL2*, *F5H*, *CCR1*, *C4H*, *C3’H*, and *CCoAOMT1* and their expression was also significantly higher as compared to wild-type plant (Figure 5E). This gene expression data suggested that there could be cross-regulation of cell wall acetylation and lignin biosynthesis pathway in *At*GELP53 OE lines. To understand more about its regulation and sensing in transgenic plants, the expression of cell wall receptor genes involved in cell wall integrity sensing were tested. Wall-associated kinases (WAKs) are known receptor-like kinases (RLKs) that function as cell wall integrity receptors and mainly recognize and bind to pectin fragments (Decreux & Messiaen, 2005). The expression of WAKs, i.e., *WAK1*, *WAK2*, *WAK3*, *WAK4*, *WAK5*, and *WAKL2*, were evaluated in *At*GELP53 OE lines, and surprisingly WAKs expression was found reduced in leaf whereas their expression was significantly upregulated in stem (Figure 5F and 5G). *THESUS1* (*THE1*) and *FERONIA* (*FER*) are also RLKs and function as sensors for various cell wall defects (Cheung & Wu, 2011). The expression of *THE1* and *FER* were significantly upregulated in both leaf and stem of *At*GELP53 OE lines (Figure 5F and 5G). To further understand about the elicitor molecules or oligosaccharides that might be interacting with wall integrity receptors or involved in generation of elicitors, we checked the expression of some wall hydrolases that might be involved in oligosaccharide release endogenously such as xyloglucan endotransglucosylase/hydrolase (XET/EH or XTH) which are known to have xyloglucan endotransglucosylase and/or endohydrolase activity (Rose et al., 2002). The expression of *XTH4*, *XTH24*, and *XTH18* was found to be significantly upregulated in *At*GELP53 OE lines (Figure 5H-J). Altogether, altered expression of cell wall acetylation genes, lignin biosynthesis genes, and more importantly cell wall integrity sensing genes explained that because of the changes in the acetylation level on individual polysaccharides in the cell wall, *At*GELP53 OE plants are sensing these changes through cell wall receptors and differentially regulating expression of cell wall related genes. This data suggested the activation of plant cell wall integrity mechanism in *At*GELP53 overexpressing lines.

**Figure 5.**
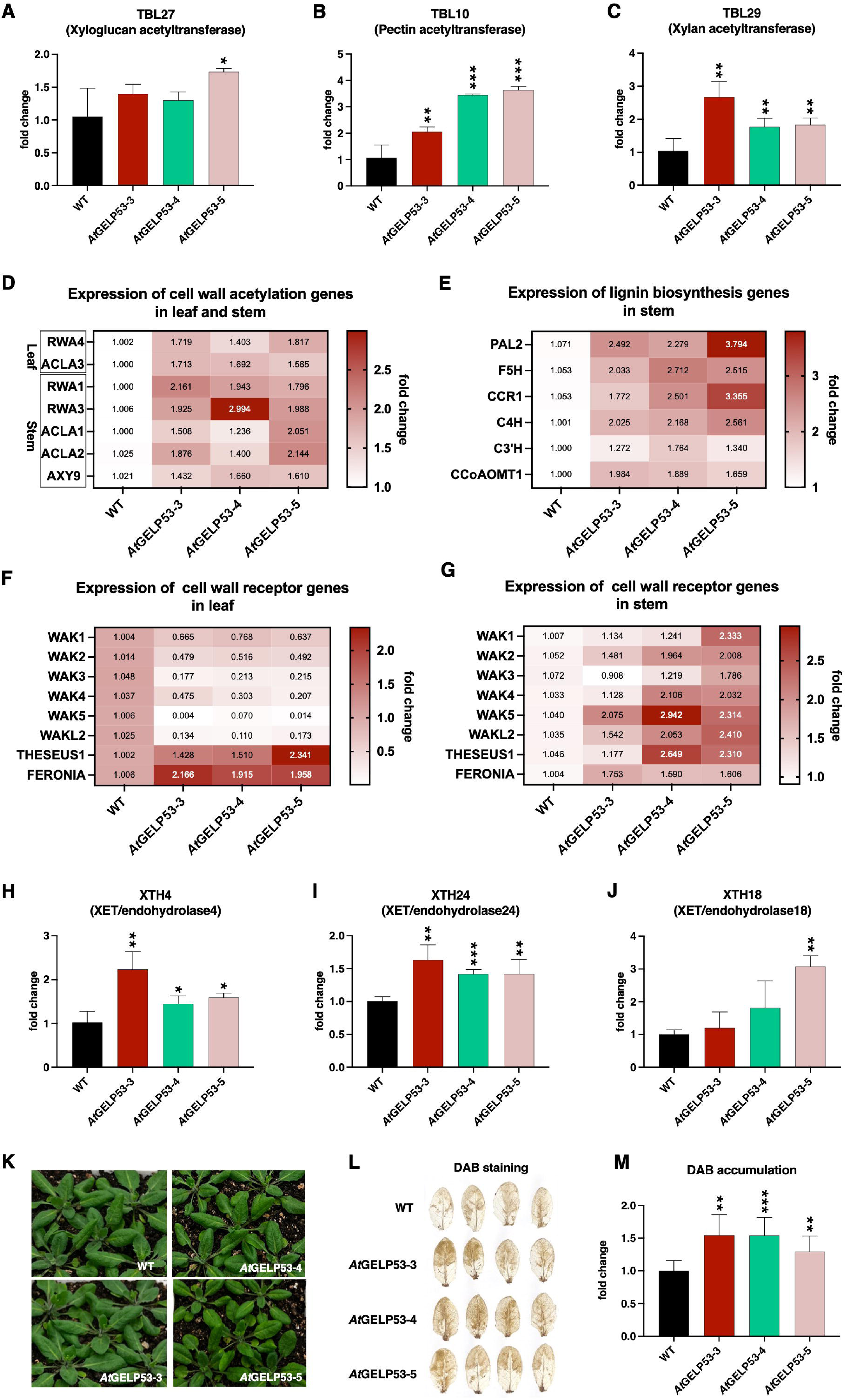
*At*GELP53 overexpression shows altered cell wall-associated gene expression and increased ROS production. **(A-C)** Expression of specific polysaccharide acetyltransferase genes such as *TBL27*, a xyloglucan acetyltransferase **(A)**, *TBL10*, a pectin acetyltransferase **(B)**, and *TBL29*, a xylan acetyltransferase **(C)** by qRT-PCR in wild-type and *35S::AtGELP53* OE lines. *ACTIN* was used as reference gene. Data represents mean ± SE, *n* = 3-4 biological replicates, Student’s t-test at \*\*\*\**p* ≤ 0.001, \*\*\**p* ≤ 0.01, \*\**p* ≤ 0.05, * *p* ≤ 0.1. **(D-E)** Heatmaps representing the expression of key genes involved in cell wall acetylation pathway such as *RWAs*, *ACLAs*, and *AXY9* in both leaf and stem **(D)**, lignin biosynthesis pathway such as *PAL2*, *F5H*, *CCR1*, *C4H*, *C3’H*, and *CCoAOMT1* in stem **(E)**, cell wall integrity sensing such as *WAKs*, *THESEUS*, and *FERONIA* in leaf **(F)** and stem **(G)** of wild-type and *35S::AtGELP53* OE lines. *ACTIN* was used as reference gene. Data represents mean ± SE, *n* = 3-4 biological replicates, Student’s t-test at \*\*\*\**p* ≤ 0.001, \*\*\**p* ≤ 0.01, \*\**p* ≤ 0.05, * *p* ≤ 0.1. **(H-J)** Expression of xyloglucan endotransglucosylase/hydrolase (XET/EH) i.e., *XTH4* **(H)**, and *XTH24* **(I)**, and *XTH18* **(J)** genes by qRT-PCR*. ACTIN* was used as reference gene. Data represents mean ± SE, *n* = 3-4 biological replicates, Student’s t-test at \*\*\**p* ≤ 0.01 \*\**p* ≤ 0.05, * *p* ≤ 0.1. **(K-M)** The rosette leaves were collected from wild-type and *35S::AtGELP53* OE lines **(K)**, H_2_O_2_ accumulation was measured by DAB staining **(L)**, and fold change of DAB accumulation **(M)** was calculated. Data represents mean ± SE, *n* = 3-4 biological replicates, Student’s t-test at \*\*\*\**p* ≤ 0.001, \*\*\**p* ≤ 0.01, \*\**p* ≤ 0.05, * *p* ≤ 0.1.

### Xyloglucanase-digested oligosaccharides derived from *At*GELP53 OE lines act as elicitors, activate expression of defence marker, integrity receptors and wall acetylation genes and induce ROS accumulation which correlates with proteomics data

There are multiple pieces of evidence suggesting that plant-derived elicitors can act as signalling molecules and can play a role in the remodelling of cell walls through cell wall-localized endogenous enzymes (Benedetti et al., 2015; Souza et al., 2017). Xyloglucan-derived elicitors can act as damage-associated molecular patterns (DAMPs) and also induce immunity in Arabidopsis and grapes (Claverie et al., 2018). Also, a previous report in tobacco showed that xyloglucan oligosaccharide (XGO) treatment alters the expression of genes associated with cell wall metabolism, transcriptional control, cell division, stress response, and signaling (Gonzalez-Perez et al., 2014). Our recent results showed that xylobiose treatment can remodel the cell wall, induce plant immunity and affect the hormone response (Dewangan et al., 2023). Since *At*GELP53 overexpressing lines exhibited less xyloglucan acetylation and remodelling in the plants, we wanted to test whether this is because of xyloglucan-derived endogenous elicitor generated by cell wall localised enzymes.

To test this elicitor-based hypothesis, we carried out mass spectrometry to compare the global proteome of wild-type and *At*GELP53 overexpressing plants. Label-free quantification identified 379 proteins common to both wild-type and *At*GELP53-5 OE lines. Of these, 108 proteins were found to be upregulated and 5 proteins were downregulated in *At*GELP5-5 OE line and 144 proteins were found uniquely in *At*GELP53-5 OE line (Figure S7A and S7B). The principal component analysis (PCA) of common proteins clearly separated the wild-type and *At*GELP53-5 overexpression line into two different groups with maximum variation of 48.2% at PC1 and 26.1% at PC2 (Figure S7C). Heatmap shows the top 50 differentially regulated proteins between wild-type and *At*GELP53-5 OE line (Figure S7D). Pathway enrichment using over representation analysis (ORA) identified pathways such as biosynthesis of secondary metabolites, MAPK signaling pathway, oxidative stress, and glutathione metabolism (Figure S7E-G). The upregulated processes such as MAPK signaling, oxidative stress and its scavenging by glutathione metabolism in *At*GELP53 overexpressing line suggesting that there might be generation of reactive oxygen species (ROS) which is the first response during elicitor-mediated signalling (Denoux et al., 2008; Dewangan et al., 2023). DAB accumulation was found higher in *At*GELP53 OE lines that correlated with our proteomics data which showed upregulation of oxidative stress proteins (Figure 5K-M).

To investigate further regarding the elicitor-mediated signaling, we extracted the cell walls of wild-type and *At*GELP53-5 overexpression plant and digested them with xyloglucanase enzyme. The xyloglucanase-digested oligosaccharides from wild-type and transgenic plants were infiltrated in rosette leaves of three-week-old Arabidopsis plants and observed ROS response by measuring DAB accumulation after 30 min and 6 h of treatment (Figure 6A). DAB staining of leaves treated with xyloglucanase-digested oligosaccharides derived from *At*GELP53-5 OE lines showed higher accumulation of ROS after 30 min as compared to the leaves treated with xyloglucanase-digested oligosaccharides derived from wild-type plant, whereas ROS accumulation was faded after 6 h of treatment (Figure 6B). We also checked the expression of some defence-related marker genes such as *PATHOGEN-RELATED1* (*PR1*), *PHYTOALEXIN DEFICIENT3* (*PAD3*), *FLG22-INDUCED RECEPTOR-LIKE KINASE1* (*FRK1*), and *WRKY33* and their expressions were also upregulated after 30 min (Figure 6C). The genes related to cell wall integrity sensing such as *THESUS1*, *FERONIA*, *WAK3* and *WAK4* were also upregulated after 30 min (Figure 6D). Interestingly, the expression of cell wall acetylation genes such as *RWA1*, *RWA2*, *RWA3*, *AXY9*, *TBL10*, and *TBL27* were also upregulated after 30 min (Figure 6E). We further performed label free proteomics studies on 30 min xyloglucan-rich derived elicitor-treated Arabidopsis leaves. Out of 519 common proteins, 161 proteins were upregulated and 122 proteins were downregulated and 82 proteins were found uniquely in leaves infiltrated with xyloglucanase-digested oligosaccharides derived from *At*GELP53-5 OE line (Figure 6F and 6G). The PCA plot of common proteins clearly separated the two groups with a maximum variation of 60.8% at PC1 and 12.8% at PC2 (Figure 6H). We also made a heatmap showing the top 50 differentially regulated proteins in these infiltrated samples (Figure S8A). The pathway analysis showed similar signalling pathways and processes such as MAPK signalling pathway, oxidative stress, and glutathione metabolism altered as found in the *At*GELP53-5 overexpression line (Figure S8B and Figure 6I and 6J). This data further confirmed the activation of elicitor-mediated signalling mainly through xyloglucan oligosaccharides in *At*GELP53 overexpressing lines.

**Figure 6.**
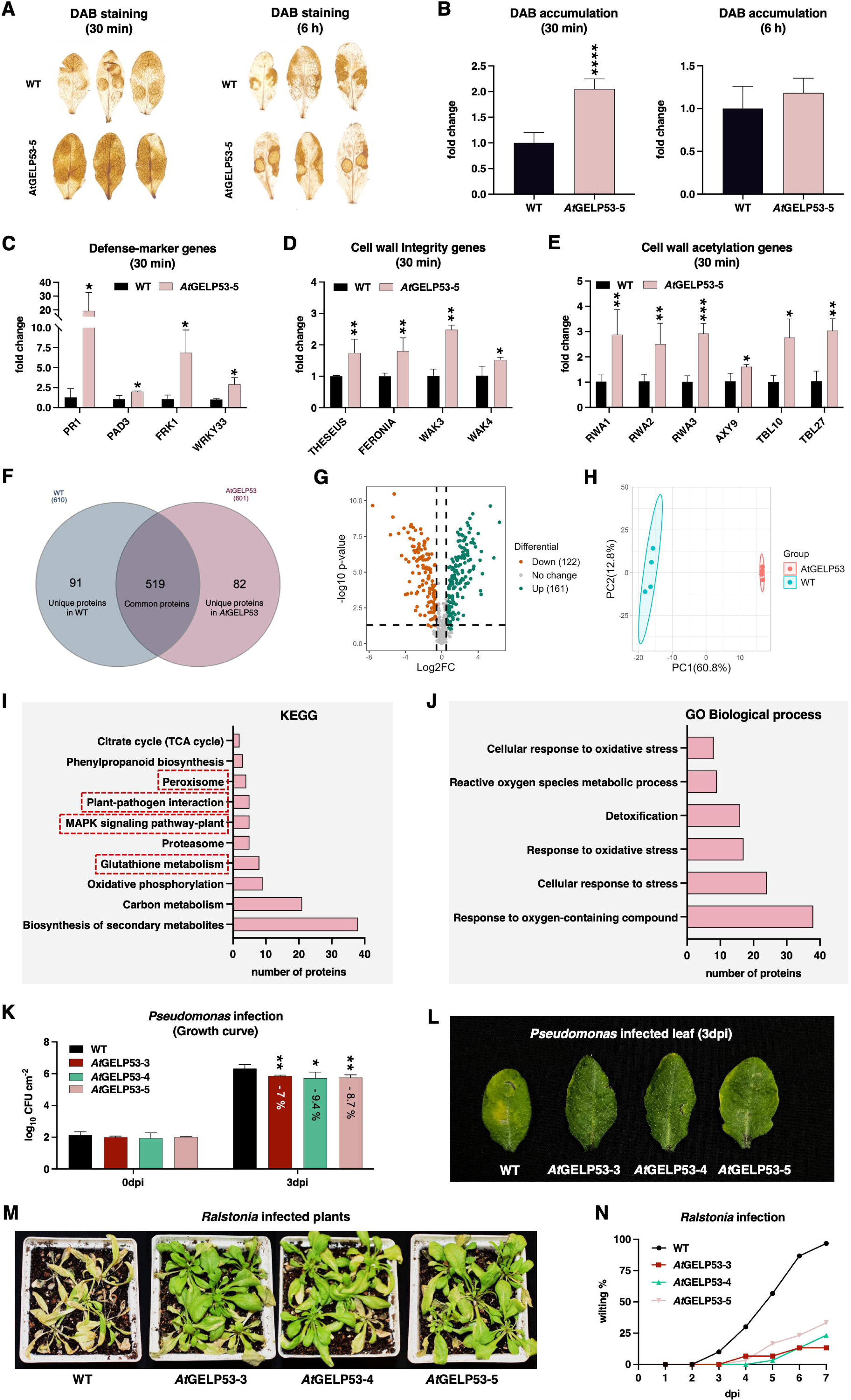
Xyloglucan-derived oligosaccharides induce elicitor-mediated responses and improve pathogen resistance. **(A)** Xyloglucanase-digested oligosaccharides derived from leaf AIR of wild-type and *At*GELP53-5 OE line were infiltrated in rosette leaves of wild-type plants and after 30 min and 6 h post infiltration, H_2_O_2_ accumulation was detected by DAB staining. **(B)** Fold change representation of DAB accumulation after 30 min and 6 h post infiltration. Data represents mean ± SE, *n* = 3-4 biological replicates, Student’s t-test at \*\*\*\**p* ≤ 0.001, \*\*\**p* ≤ 0.01, \*\**p* ≤ 0.05, * *p* ≤ 0.1. **(C-E)** The expression of defense-marker genes such as *PR1*, *PAD3*, *FRK1*, and *WRKY33* **(C)**, cell wall integrity genes such as *THESEUS*, *FERONIA*, *WAK3*, and *WAK4* **(D)**, and cell wall acetylation such as *RWA1*, *RWA2*, *RWA3*, *AXY9*, *TBL10*, and *TBL27* **(E)** was analyzed in leaf samples after 30 min post infiltration by qRT-PCR. *GAPDH* was used as reference gene. Data represents mean ± SE, *n* = 3-4 biological replicates, Student’s t-test at \*\*\*\**p* ≤ 0.001, \*\*\**p* ≤ 0.01, \*\**p* ≤ 0.05, * *p* ≤ 0.1. **(F)** Venn diagram generated by InteractiVenn software depicting the number of common and unique proteins expressed after 30 min post infiltration. **(G)** Volcano plot depicting the number of upregulated and downregulated proteins in leaves infiltrated with xyloglucanase-digested oligosaccharides derived from *At*GELP53-5 OE line after 30 min post infiltration. **(H)** Principal component analysis (PCA) plot shows a clear separation of leaf samples infiltrated with xyloglucanase-digested oligosaccharides derived from *At*GELP53-5 OE line from WT samples. **(I-J)** Pathway analysis of upregulated and unique proteins identified in leaves infiltrated with xyloglucanase-digested oligosaccharides derived from *At*GELP53-5 OE line after 30 min post infiltration by ShinyGO (v0.80) tool; selected pathways from KEGG involved in elicitor-mediated signaling **(I)** and representation of altered biological processes by Gene Ontology (GO); Biological process **(J)**. **(K-L)** Growth of *Pseudomonas syringae pv. tomato DC3000* (*PstDC3000*) in wild-type and *35S::AtGELP53* OE lines at 0dpi and 3dpi **(K)**, and a picture of plants showing disease symptoms as yellow patches at 3dpi **(L)**. dpi; days post infiltration. Data represents mean ± SE, *n* = 3-4 biological replicates, Student’s t-test at \*\*\*\**p* ≤ 0.001, \*\*\**p* ≤ 0.01, \*\**p* ≤ 0.05, * *p* ≤ 0.1. **(M-N)** Picture of wild-type and *35S::AtGELP53* OE plants infected with *Ralstonia solanacearum F1C1* **(M)** and the percentage of wilted plants was plotted as a line chart **(N)**. A minimum of 10 Arabidopsis plants were infected in each experiment, and the experiment was independently repeated three times. Data represents mean ± SE, *n* = 15 biological replicates and the experiment was repeated three times with similar results.

### *At*GELP53 OE plants show resistance against *Pseudomonas syringae* and *Ralstonia solanacearum*

Modifications in the plant cell wall have been shown to impact plant responses to biotic stress. The cell wall changes could induce cell wall-mediated immunity (CWI) in plants and trigger disease resistance responses (Bacete et al., 2018; Molina et al., 2024; Molina et al., 2021; Yu et al., 2024). To study such responses in *At*GELP53 overexpression plants due to their altered cell wall structure and defense response, we challenged *At*GELP53 OE lines with plant pathogens such as *Pseudomonas syringae* and *Ralstonia solanacearum*. An equal number of *P. syringae* colonies were observed at day 0, whereas at day 3, *P. syringae* growth was significantly reduced in *At*GELP53 OE lines as compared to wild-type plants and disease symptoms, i.e., yellow patches were not much apparently visible on leaves of transgenic plants (Figure 6K and 6L). *Ralstonia solanacearum* which infects plants through their roots, colonizes the xylem vessels and cause bacterial wilt disease in plants. Apparently, total acetylation was lower in root tissue of *At*GELP53 lines. Therefore, we treated roots of wild-type and *At*GELP53 OE plants with *Ralstonia solanacearum* strain *F1C1* at a density of 1×10^8^ CFU/ml and observed wilting of plants (Figure 6M). The wilting symptoms were recorded from day 1 to day 7 of treatment. The transgenic *At*GELP53 OE plants showed less than 25% wilting after 7 days of infection compared to wild-type plants which were completely wilted after 7 days (Figure 6N). Together, these pathogen infection assays showed that *At*GELP53 overexpression plants were resistant to *Pseudomonas syringae* and *Ralstonia solanacearum* infection.

## DISCUSSION

Plant cell wall *O*-polysaccharide acetylation needs to be balanced because it has a significant role in maintaining the overall cell wall architecture to sustain growth and development. The maintenance and balancing of wall acetylation are done through polysaccharide acetyl transferases and esterases. Various Golgi-localized non-specific and specific polysaccharide acetyl transferases have been studied extensively in plants, but very few polysaccharide esterases having antagonistic functions are identified and characterized. So far, pectin acetyl esterases (PAE) from carbohydrate esterase 13 (CE13) family and acetyl xylan esterases (AXE) from GELP family are known in plants (de Souza et al., 2014; Gou et al., 2012; Orfila et al., 2012; Rastogi et al., 2022; Zhang et al., 2017). There is no report of xyloglucan acetyl esterase in plants. In our study, we have functionally characterised *At*GELP53 as xyloglucan acetyl esterase from the GELP family and its role in elicitor-mediated induction in plant immunity.

The Arabidopsis GELP family has 105 protein members that include both esterases and lipases, and so far, most of the lipases and very few esterases are characterized as listed in the figure (Figure 1A and 1B). In summary, GELP family members show diverse functions; and impact processes including pollination, disease resistance via ethylene and auxin metabolism, cutin biosynthesis, glucosinolate metabolism, and fatty acid biosynthesis in different tissue types (Agee et al., 2010; Chen et al., 2012; Hong et al., 2017; Huang et al., 2015; Jancowski et al., 2014; Lee et al., 2009; Mayfield et al., 2001; Nagano et al., 2008; Naranjo et al., 2006; Oh et al., 2005; Takahashi et al., 2010; Updegraff et al., 2009; Zhang et al., 2006).

In 2002, *AtFXG1/AtGELP33* was reported to have fucosidase activity in the cell wall, thereby showing the role of GELP family in cell wall modification (De La Torre et al., 2002). *At*AXE1/*At*GELP7 is characterized as plasma-membrane localized acetyl xylan esterase (AXE) that specifically deacetylates xylan, and its overexpression improves biomass digestibility (Rastogi et al., 2022). Here, *At*GELP53 from this study shares the same clade Id of the GELP family and shows 70% sequence similarity with *At*AXE1/*At*GELP7 (Figure 1B). Our transient and stable GFP expression studies revealed that *At*GELP53 is also localized in plasma membrane and has esterase activity (Figure 1C-F). Previously identified as belonging to the GELP family, acetyl xylan esterase from rice is localized in the Golgi membrane, and it plays a role in de-esterification during synthesis (Zhang et al., 2017). However, *At*GELP53 is functioning post-synthesis of the cell wall. *At*GELP53 promoter is expressed in all the tissue but predominantly found in leaf primordia, root tip and reproductive organ as compared to mature stem or leaf tissue (Figure 2A). Although the acetyl content was reduced in all tissues of *At*GELP53 overexpressing lines, this can normally function as esterase in primary cell wall tissue (Figure 2J-L). However, no aberrant growth phenotype was observed in overexpressing GELP53 lines, on the contrary, plant showed enhanced overall height and root length (Figure 2B and 2C and Figure 2H and 2I). This could be because 8.4% or 13.6% reduction in acetylation is tolerated by plants (Figure 2J and 2L). Another possible explanation is that xyloglucan deacetylation is dispensable for plants or xyloglucan deacetylation is compensated by more xylan and pectin acetylation which can make the cell wall stronger (Gille et al., 2011; Pawar et al., 2013). Cell type-specific probing of polysaccharide acetylation can address the role of acetylated polysaccharides in normal growth conditions.

*At*GELP53 overexpressing lines showed a reduction in acetyl content in leaf, root, and stem tissue. Xyloglucan structure is different in tissue types in the same plants in terms of substitution pattern, making it complex to interpret xyloglucan fingerprinting data after xyloglucanase digestion (Schultink et al., 2014). Probing of acetylation in different fractions of polysaccharides revealed that it is because of xyloglucan deacetylation (Figure 3A-D). Our data indicated less accumulation of acetylated FLXG and FXXG (Figure 3G and 3H) after xyloglucanase digestion which was also observed in *tbl27* or *axy4* (Gille et al., 2011). Also, the xyloglucan-derived non-acetylated XXXG oligosaccharides were detected more in *At*GELP53 overexpression lines further correlated with reduction in acetylation of xyloglucan-rich fraction (Figure 3E and 3F). Moreover, LM15 antibody which recognizes XXXG, its signals were detected more in transgenic lines as compared to wild-type (Figure 4A). The structural model of *At*GELP53 showed the typical fold of SGNH hydrolase family with conserved catalytic triad residues (Ser40, Asp344, and His347). Additionally, *At*GELP53 contains six possible disulfide bridges (Cys28-Cys374, Cys122-Cys132, Cys154-Cys161, Cys229-Cys251, Cys311-Cys334C, and Cys312-Cys326) and four potential N-linked glycosylation sites (Asn136, Asn319, Asn371, and Asn382) (Figure 3K). Interestingly, the disulfide bridge loop Cys311-Cys334 likely regulates the substrate binding to the active site cleft, as shown in LPLA. Our docking analysis indicated that the large-size active site cleft of *At*GELP53 can accommodate a xyloglucan substrate, corroborating with findings from the current study that *At*GELP53 could act as a xyloglucan acetyl esterase (Figure 3M).

The altered pattern of pectin and xylan was further confirmed through enzymatic oligosaccharide profiling and immunolocalization studies. Polygalacturonase digestion can be primarily inhibited by the presence of pectin methyl ester (Jermendi et al., 2022). HGs are *O*-acetylated, whether they hinder polygalacturonase activity is not yet clear. However, a reduction in pectin acetylation was observed after overexpression of poplar pectin acetyl esterase in *Nicotiana benthamiana*, and the pectin of *Pt*PAE-OE line was less digestible (Gou et al., 2012)*. At*GELP53 overexpressing lines showed higher acetyl/GalA after pectate lyase digestion and an increase in the release of non-methylated GalA3 and GalAMe by polygalacturonase (Figure 3B and 3I). This suggests that pectin acetylation and/or methylation pattern is affected in these *At*GELP53 overexpression plants which can be explored in future by NMR and fingerprinting methods since both substitutions can impact cell wall remodeling (Shahin et al., 2023). *At*GELP53 lines also showed an increase in the acetylation of the xylan-rich fraction and less abundant shorter-chain XOS and more abundant longer-chain XOS after xylanase digestion (Figure 3D and 3J). Several of our previous reports in Arabidopsis and Poplar demonstrated that *in planta* reduction in xylan acetylation increases xylan digestibility by xylanases (Pawar et al., 2017; Pawar et al., 2016; Rastogi et al., 2022). And in monocots, more xylan acetylation in rice *bs1* mutant decreases xylan digestibility (Zhang et al., 2017). Overall, this data further validates that xylan acetylation hinders xylanase accessibility and it is a promising target to improve cell wall properties.

Polysaccharide extractability depends on crosslinking between polymers via acetyl substitutions on the wall polysaccharides (Kong et al., 1992). The more extractability of polysaccharides could result from less crosslinking between wall polymers due to the presence of less decorations on polysaccharides. In contrast, less extractability of polysaccharides could result from more crosslinking between wall polymers due to more decorations on polysaccharides. The polysaccharide extractability assays such as ELISA-based glycome profiling of extracted wall polysaccharides showed that xyloglucan extractability is higher in 4MKOH extract as compared to wild-type plants, which could be due to less acetyl decorations on xyloglucan in transgenic lines whereas pectin extractability is low in CE, 1MKOHE, and 4MKOHE and xylan extractability is also low in 1MKOHE, and 4MKOHE, which could be due to higher acetyl decorations on pectin and xylan (Figure 4B and 4C). This extractability data nicely correlates with the observed acetylation levels on different polysaccharides in the *At*GELP53 transgenic lines. We have a limited understanding of how acetylation influences polysaccharide extractability, but this data indicates less acetylation can lead to more polysaccharide extractability. This can be further tested in different acetylated mutants to confirm our observations.

Xylan *O*-acetylation and lignin content in the cell wall are the major determining factors that affect cellulose digestibility. In our previous study of *At*AXE1/*At*GELP7, xylan acetylation and lignin content was reduced upon *At*AXE1 overexpression and saccharification efficiency was enhanced (Rastogi et al., 2022). Interestingly, here in *At*GELP53 overexpression plants, we observed the exact opposite scenario where xylan has more acetylation and lignin content is also higher and, saccharification efficiency was reduced in transgenic lines (Figure 3D, 4E, 4H and 4I).

Acetylation changes in wall polysaccharides seem to be found parallel to the changes in cell wall acetylation genes in *At*GELP53 overexpression lines. Upregulation of RWA, AXY9, TBL10, and TBL29 correlating with the increased acetylation on pectin and xylan suggests a tight acetylation regulation in the cell wall (Figure 5A-D). The lignin biosynthetic machinery is also upregulated, leading to increased lignin content upon *At*GELP53 overexpression (Figure 5E). Further, the upregulation of cell wall integrity receptors like WAKs, THESEUS, and FERONIA represent change in plant cell wall integrity signalling due to *At*GELP53 overexpression which can alter the cell wall structure which was also detected by ROS accumulation (Figure 5F, 5G and 5M). We found enhanced expression of xyloglucan endotransglucosylase/hydrolases (XET/XTH) in *At*GELP53 OE lines which could alter oligosaccharide profile in apoplastic space that can act as potential elicitors (Anderson & Kieber, 2020; Rose et al., 2002) (Figure 5H-J). From previous studies, it is well known that cell wall-derived oligosaccharides, especially from cellulose, pectin, xylan, and xyloglucan act as endogenous elicitors known as DAMPs which can bind to plasma-membrane localized wall integrity receptors, activate MAPK pathway for downstream signaling and also trigger ROS accumulation (Benedetti et al., 2015; Claverie et al., 2018; Dewangan et al., 2023; Souza et al., 2017). MAPK pathway and ROS accumulation influence the downstream signalling and, through transcriptional reprogramming, induce a variety of responses in plants. Likewise, in our case, we saw an alteration in the cell wall acetylation genes that compensated for the loss of acetylation in the cell wall, higher lignin content and ROS production. This was confirmed through elicitor-based assay where xyloglucanase-digested oligosaccharides derived from *At*GELP53 overexpression lines when infiltrated in wild-type leaf, induced ROS accumulation and activated defence-marker, wall integrity and cell wall acetylation genes after 30 min of treatment (Figure 6A-E). Such type of PTI response was also observed in grapevine after treatment with mixture of xyloglucan-derived oligosaccharides (Claverie et al., 2018). Furthermore, our proteomics data identified proteins that are involved in oxidative stress and MAPK signaling pathway upon *A*tGELP53 overexpression and in 30 min treated leaf samples with xyloglucanase-digested oligosaccharides derived from *At*GELP53 overexpression lines (Figure S7E-G and Figure 6I and J). The *At*GELP53 overexpression plants also showed resistance against *Pseudomonas syringae* and *Ralstonia solanacearum* due to changes in the cell wall structure and/or DAMP-derived induction in immunity that can provide resistance to fight the disease (Figure 6K-N). Although the mechanism of altered defence regulation depends on type of cell wall modification, crop and pathogen (Molina et al., 2024), the role of GELP53 in improving plant immunity can be explored further in different crop species. These findings suggest that upon *A*tGELP53 overexpression due to loss of xyloglucan acetylation, there may be the activation of oligosaccharide-based elicitor-mediated signaling through wall integrity receptors that may further translate signal internally via MAPK pathway or other unknown pathway and reprogram the transcriptional machinery by altering the expression of cell wall acetylation and lignin biosynthetic genes.

In summary, we have characterized the role of *At*GELP53 in xyloglucan acetylation, and its overexpression alters cell wall composition (lignin, polysaccharide acetylation) and induces plant defence. We proposed a possible mechanism of how deacetylation of xyloglucan can alter oligosaccharide-mediated signalling by maintaining cell wall integrity response which is summarised in the schematic model (Figure 7).

**Figure 7.**
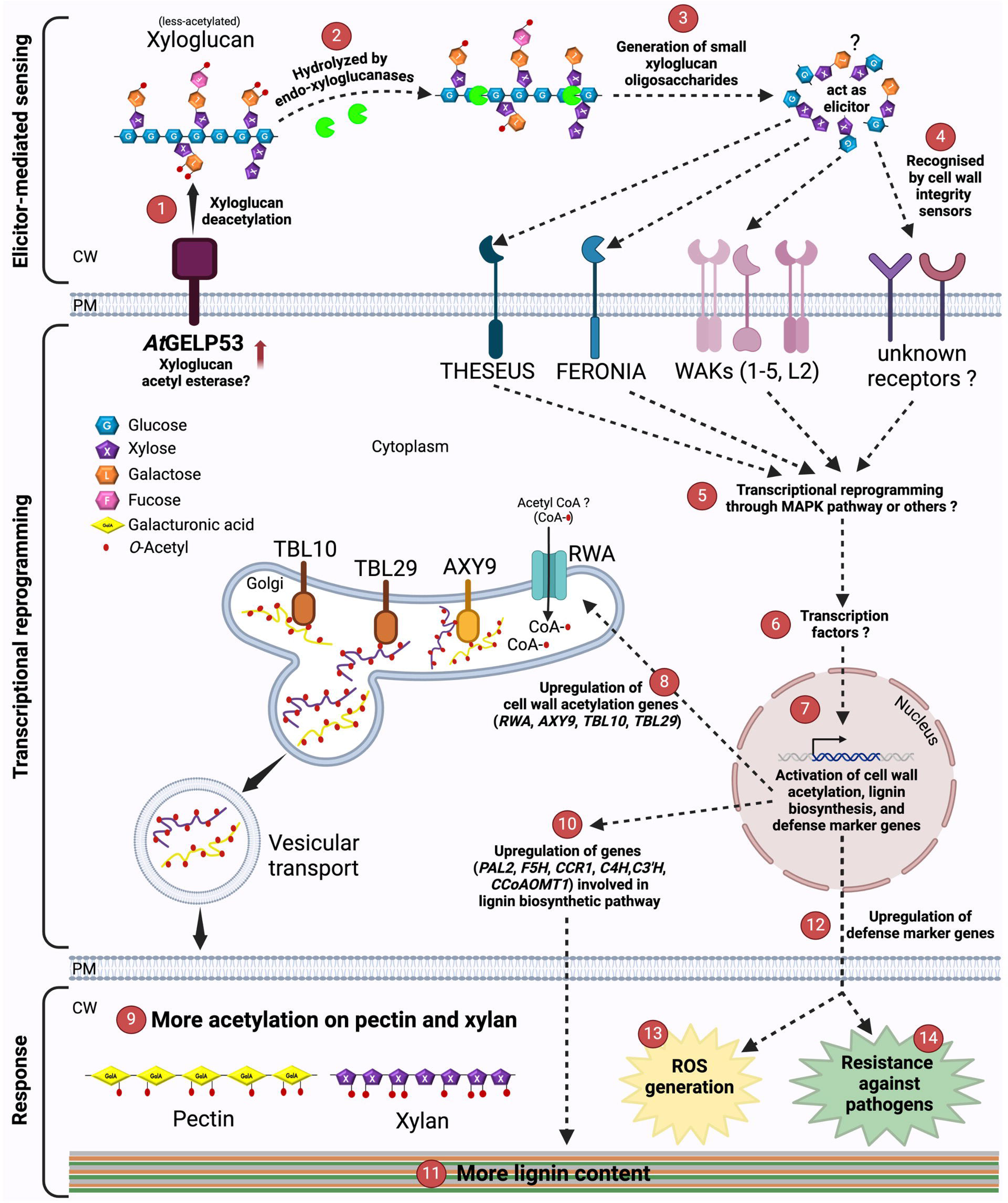
Schematic representation of elicitor-mediated sensing, transcriptional reprogramming, and response upon *At*GELP53 overexpression in Arabidopsis. **(1)** *At*GELP53 protein is located in the plasma membrane and its overexpression decreases xyloglucan acetylation. **(2)** Deacetylation of xyloglucan in the cell wall may increase accessibility for xyloglucanases or other hydrolytic enzymes. **(3)** Either more or unique xyloglucan fragments may be generated which can act as endogenous elicitors. **(4)** Xyloglucan-derived elicitors may be recognized by cell wall integrity and sensing receptors such as THESEUS, FERONIA, WAKs (1-5, L2) or some unknown receptors. **(5)** Interaction of xyloglucan-derived elicitor molecules with cell surface integrity receptors initiates signaling and transcriptional reprogramming through MAPK pathway or some other pathway. **(6)** This signaling further initiates downstream signaling through some transcription factors and reprograms the transcriptional machinery inside the cell which can **(7)** lead to the upregulation of cell wall acetylation, lignin biosynthesis, and defense-marker genes. **(8)** Elicitor-mediated signaling led to the upregulation of cell wall acetylation genes such as *RWA, AXY9, TBL10,* and *TBL29* that are localized in Golgi membrane and increases acetylation mainly on pectin and xylan polysaccharides. **(9)** Hyper-acetylated pectin and xylan polysaccharides from the Golgi lumen are transported and deposited into the cell wall and maintain the required amount of polysaccharide *O*-acetylation in the cell wall. **(10)** & **(11)** Upregulation of lignin biosynthetic genes lead to an increase in lignin content in the cell wall. **(12)** Upregulation of defense-marker proteins induces cell wall-mediated plant immunity. **(13)** The generation of reactive oxygen species (ROS) is because of elicitor-mediated signaling. **(14)** The alteration in cell wall structure, ROS accumulation, and induction in plant immunity may have provided resistance to *At*GELP53 overexpression plants against pathogens.

## METHODS

### Generation of phylogenetic tree

Gene IDs of the Arabidopsis GELP family were taken from (Lai et al., 2017). Amino acid sequences of 105 *At*GELPs were fetched from plantgenie.org, and the phylogenetic tree was created by MEGA11 software using the maximum likelihood (ML) method with 1000 bootstraps.

### Plant growth conditions

*Arabidopsis thaliana* (*Columbia-0*) and *Nicotiana benthamiana* were grown in a walk-in plant growth chamber at 22°C under 16h light/8h dark cycle conditions. These plants were used in all experiments.

### *AtGELP53* gene cloning

Using *AtGELP53* (*AT2G27360*) gene-specific forward (PCWL37) and reverse (PCWL38) primers, this gene was amplified from Arabidopsis leaf cDNA and cloned first into entry vector pDONR^TM^ 207 and then into destination vectors i.e., pCC0995 (containing constitutive 35S promoter), pSITE-2CA (containing N-terminal GFP tag), and pKGWFS7 (containing GUS) by gateway cloning method using Gateway BP Clonase II enzyme mix (11789, Invitrogen, Canada) and Gateway LR Clonase II enzyme mix (11791, Invitrogen, Canada). After transformation into DH5alpha *E.coli* competent cells, the confirmed clones were transformed into *Agrobacterium tumefaciens* GV3101 strain.

### Subcellular localization

Agrobacterium cells containing *35S::GFP-AtGELP53* were grown at 28°C and resuspended in infiltration medium containing 10mM MES buffer (RM1128, HiMedia, India) and 10mM MgCl_2_ (MB237, HiMedia, India) pH-5.6 with 0.4 OD and induced with 100 µM acetosyringone (RM9145, HiMedia, India) for at least 4 hours and infiltrated in Nicotiana leaves. On 3^rd^ day post-infiltration, the leaves were visualised under LEICA SP8 confocal microscope in a 40X oil immersion lens.

### Generation of *Arabidopsis thaliana* transgenic lines

Agrobacterium culture having *35S::AtGELP53* were resuspended in transformation medium having 5% sucrose (GRM3063, HiMedia, India) and 0.05% silwet-77 (PCT1554, HiMedia, India). Further procedure for floral dip transformation was done according to the protocol explained in (Clough & Bent, 1998). Dried seeds were collected and successfully transformed plants were selected by spraying glufosinate ammonium (BASTA) (C45520, Sigma-Aldrich, Switzerland). T3 homozygous lines were used for all the experiments.

### Esterase and lipase activity

Crude proteins were extracted from plant tissue and esterase activity was performed with p-Nitrophenyl Acetate (18432, SRL, India) substrate, whereas lipase activity was performed with p-Nitrophenyl Palmitate (N2752, Sigma-Aldrich, Mongolia) substrate as described in (Rastogi et al., 2022).

### Protoplast isolation

Protoplasts from leaves of stable *GFP-AtGELP53* Arabidopsis lines were isolated by the Tape-Arabidopsis Sandwich method as described in (Wu et al., 2009). The isolated protoplasts were visualised under LEICA SP8 confocal microscope in a 40X oil immersion lens.

### Genotyping of T-DNA mutant

From Arabidopsis Biological Resource Centre (ABRC), SALK_078877C line of *AtGELP53* was obtained and genotyping was done using left primer-LP(PCWL15), right primer-RP(PCWL16), and T-DNA specific border primer-BP (PCWL29) (Table S3) to select homozygous lines of mutant *atgelp53*.

### GUS assay

GUS staining solution was prepared by dissolving X-Glucurono sodium salt (MB089, HiMedia, India) in DMSO, 50mM sodium phosphate buffer, 100mM potassium ferricyanide (GRM1034, HiMedia, India), 100mM potassium ferrocyanide trihydrate (GRM1048, HiMedia, India), and 0.1% triton. Different tissues of stable *pAtGELP53::GUS* transgenic lines were dipped in GUS staining solution and incubated overnight at 37°C. For destaining, ethanol (3):acetic acid (1) was used and incubated until decolorised. Images were captured under a light microscope.

### RNA isolation, cDNA synthesis, and qRT-PCR

Total RNA was extracted from grounded leaf and stem tissues of Arabidopsis using Trizol method (15596018, Invitrogen, Canada). cDNA was synthesized from 1ug RNA using iScript cDNA Synthesis kit (1708891, BioRad, USA). For qRT-PCR, *ACTIN2* (*AT3G18780*) and GAPDH (*AT1G13440*) were used as reference genes. The expression of *AtGELP53* and other cell wall-related genes was measured using gene-specific qPCR primers listed in Table S3. The fold change of each gene with respect to reference gene was calculated by ΔΔCt method.

### Seed germination on MS media

Seeds were sterilised in 70% ethanol followed by 100% ethanol and germinated on half MS plates prepared by dissolving MS salt (Murashige & Skoog medium w/ CaCl_2_, vitamins & sucrose; w/o agar, PT099, HiMedia, India) in miliQ water, pH was set to 5.8 and solidified using 0.8% phytoagar (PCT0901 HiMedia, India). The plates were initially kept in dark at 4°C for two days and then transferred to light in a plant growth chamber. The root lengths of 12-day-old seedlings were measured by ImageJ software.

### Alcohol Insoluble Residue (AIR) preparation

Fresh tissue grounded in liquid nitrogen and dried tissue grounded by Qiagen TissueLyser II was treated with 80% ethanol and incubated at 70°C for 30 min. The pellet left after centrifugation was then treated with 70% ethanol followed by chloroform: methanol (1:1) and finally washed in acetone, and the pellet was dried.

### Cell wall acetyl content analysis

1 mg of AIR was treated with 1M NaOH for de-esterification for 2 h and neutralized with 1M HCl. Acetyl content was analyzed in the saponified supernatant by acetic acid kit (K-ACET, Megazyme, Ireland).

### Sequential extraction of cell wall polysaccharides

The sequential extraction was done according to the flow chart (Figure S4). Briefly, leaf AIR was treated with 50mM ammonium formate (50504, SRL, India) buffer for 24 h at 37°C, and the supernatant was collected as pectin-rich fraction I. The remaining pellet was then digested with pectate lyase (E-PCLYAN2, Megazyme, Ireland) at 40°C for 24 h and the supernatant was collected as pectin-rich fraction II. The pellet rich in xyloglucan and xylan was digested with xyloglucanase (E-XEGP, Megazyme, Ireland) at 50°C for 2 days, and the supernatant was collected as xyloglucan rich fraction. These fractions were freeze-dried and redissolved in water. The remaining pellet rich in xylan was washed with acetone and dried. Then, all the fractions were analysed for acetic acid (K-ACET, Megazyme, Ireland), galacturonic acid by biphenyl method, and xylose (K-XYLOSE, Megazyme, Ireland) content.

### Enzymatic oligosaccharide fingerprinting by liquid chromatography-mass spectrometry (LC-MS)

AIRs were digested with 1 U/mg dried weight of either *Pectobacterium carotovorum* endo-Polygalacturonanase (Megazyme, Bray, Ireland), *Paenibacillus sp*. xyloglucanase, or *Neocallimastix patriciarum* endo-1,4-β-Xylanase in 50mM ammonium acetate buffer (pH 5) at 37°C for 18 h. The oligosaccharides released from digestion were separated according to (Voxeur et al., 2019). Chromatographic separation was performed on an ACQUITY UPLC Protein BEH SEC Column (125Å, 1.7 μm, 4.6 522 mm × 300 mm, Waters Corporation, Milford, MA, USA) coupled with a guard Column BEH SEC Column (125Å, 1.7 μm, 4.6 mm × 30 mm). Elution was performed in 50mM ammonium formate, 0.1% formic acid at a flow rate of 0.4 ml/min, with a column oven temperature of 40°C. The injection volume was set to 10 μl. Quantitative evaluation of cell wall fragments was conducted using an HPLC system (UltiMate 3000 RS HPLC system, Thermo Scientific, Waltham, MA, USA) coupled to an Impact II Ultra-High Resolution Qq-Time-Of-Flight (UHR-QqTOF) spectrometer (Bruker Daltonics, Bremen, Germany) equipped with an electrospray ionization (ESI) source in negative mode. The end plate offset set voltage was 500 V, capillary voltage was 4000 V, nebulizer pressure was 40 psi, dry gas flow was 8 L/min, and the dry temperature was set to 180°C. The Compass 1.8 software (Bruker Daltonics) was used to acquire the data, and peak areas were integrated manually.

### Molecular modelling and docking

The three-dimensional structural model of *At*GELP53 lacking signal peptide (residues 1-24) was predicted using the AlphaFold2 server (Jumper et al., 2021) since its experimental structure is currently unavailable in the protein data bank (PDB). The *At*GELP53 model was used in molecular docking to build a possible ligand (xyloglucan) at the active site through AutoDock Vina (Eberhardt et al., 2021). A three-dimensional model of xyloglucan from *Caldicellulosiruptor lactoaceticus* GH74 (*Cl*GH74A) enzyme complex with LLG xyloglucan (PDB ID: 6P2M) was used in docking (Vieira et al., 2021). A single docked pose with the best score was used for the analysis.

### Immunolocalization of glycan epitopes

Arabidopsis main stems were cut into small pieces (1-5 mm) and fixed in chilled ethanol(3):acetic acid(1), followed by dehydration in graded ethanol series for 30 min at each step. The dehydrated tissue pieces were infiltrated with LR White (1):100% ethanol(1) for 24 h; LR White only for 24 h (three times) and solidified by heating at 80°C for 2 h. The tissue blocks were trimmed using hand-microtome to cut sections of 5 μm size. The sections were stained with Toluidine Blue O (T3260, Sigma-Aldrich, USA) and immunolabeled with different wall glycan antibodies (LM10, LM11, LM15, LM20) as explained in (Avci et al., 2012). The sections were observed under Nikon DS-Qi2 ECLIPSE T*i* fluorescence microscope.

### Glycome profiling by ELISA

Arabidopsis main stem AIR was treated with 50mM ammonium oxalate (pH 5.0) followed by 50mM sodium carbonate, 1M KOH, and 4M KOH and the supernatant collected at each step, further dialyzed and total sugar was estimated by phenol-sulfuric acid method. All the wall extracts were diluted to a sugar concentration of 500 ng/ml, 50 µl added to each well in an ELISA plate and kept for drying overnight at 37°C. ELISA was performed using different wall glycan antibodies (LM10, LM11, LM15, LM19, LM 28, CCRC M7) as per the protocol described in (Pattathil et al., 2012).

### Cell wall composition

#### Hemicellulosic monosugar composition by ion chromatography (IC)

2 mg of AIR was treated with 1.3M HCl and incubated at 100°C for 1 h. After 1 h, the samples were neutralized with 1.3M NaOH, centrifuged at 1500 g for 10 min and supernatant was collected and diluted 25 times. 100ul of diluted samples were transferred to vials and run on the Dionex system.

#### Pectin content

2 mg of AIR was treated with 2M H_2_SO_4_ at 100°C for 6 h. After neutralization with 2M NaOH, galacturonic acid content was analyzed by uronic acid kit (K-URONIC, Megazyme, Ireland).

#### Cellulose content

3 mg of AIR was treated with Updegraff reagent [acetic acid (8): nitric acid (1): water 2)] to remove hemicellulosic glucose. The remaining cellulose-rich pellet was further treated with 72% sulfuric acid, and glucose content was analyzed in hydrolysate by anthrone assay. Absorbance was taken at 600 nm (Updegraff, 1969).

#### Lignin content

Acetyl Bromide Soluble Lignin (ABSL) method was used to quantify lignin. Briefly, 1 mg of AIR was treated with freshly prepared acetyl bromide (135968, Sigma-Aldrich, Mongolia) at 50°C for 2 h, neutralized with 2M NaOH and hydroxylamine hydrochloride (159417, Sigma-Aldrich, USA), and absorbance was taken at 280 nm (Foster et al., 2010).

### Xylanase digestion

3 mg of AIR was digested with endo 1,4-ß-Xylanase (E-XYLNP, Megazyme, Ireland) at 60°C for 6 h and xylose content was analyzed by xylose kit (K-XYLOSE, Megazyme, Ireland).

### Saccharification

5 mg of AIR was incubated with cellulase enzyme blend (SAE0020, Sigma-Aldrich, USA) at 50°C for 24 h and in the supernatant glucose release was analyzed by Glucose oxidase (GOD) - Peroxidase (POD) assay (Van Acker et al., 2016).

### Proteomics analysis

Leaf tissue was grounded to fine powder and resuspended in radioimmunoprecipitation assay (RIPA) buffer (R0278, Sigma-Aldrich, USA) containing protease phosphatase inhibitors and incubated for 1 hour at 4°C. After centrifugation at 16000 rpm for 30 min, the supernatant was collected, followed by quantification by BCA (23225, Thermo Scientific, USA). 500-1000 µg of protein was taken for sample preparation which was precipitated in chilled acetone. Samples were centrifuged at 6000g for 45 minutes at 4°C and washed twice with acetone. The obtained pellet was dissolved in 8M urea followed by quantification by BCA. 50 µg of protein was processed further and incubated with dithiothreitol (DTT, 10mM) (53819, Sigma-Aldrich, USA) for 1 hour at 56°C, which was followed by incubation in dark for 1 h with iodoacetamide (IAA, 20mM) (16125, Sigma-Aldrich, USA). The protein samples were then digested with trypsin protease (90057, Thermo Scientific, USA) in 1:20 ratio at 37°C for 18 h and dried in a speed vacuum concentrator. The digested samples were redissolved in 200 µl of solution A (98% water + 2% ACN + 0.1% Formic acid) and desalted using C18 desalting cartridges (18600383, Waters, USA) which were pre-equilibrated with solution B (100% water + 0.1% Formic acid). The samples were bound to C18 bed two times and unbound fractions were washed out with 200 µl of solution A (5X). The bound peptides were eluted in 200 µl of solution C (70% ACN + 0.1% FA) and vacuum-dried. Frozen samples were reconstituted in solution A for further LC-MS/MS analysis. Each sample was acquired in triplicates on Zeno TOF 7600 (Sciex, Concord Canada) which was coupled with Luna Micro Trap (20 x 0.3 mm, 5 µm, 100 Å Phenomenex) column. The peptides were eluted from the nanoEaseTM M/Z C18 (150 mm x 300 µm, 1.8 µm, 100 Å Waters) analytical column using Acquity M class UPLC (Waters) at a flow rate of 5 µl/min in a linear gradient of 5% to 80% solvent B (100% ACN + 0.1% Formic acid) in 45 min with a total run time of 50 min. The mode of acquisition was information dependent (IDA) with a TOF/MS survey scan (300-1500 m/z) along with an accumulation time of 100 ms, and MS/MS spectra were collected for 20 ms (200-1800 m/z). MaxQuant software (v1.6, Max Planck Institute of Biochemistry, Germany) was used to identify and quantify proteins using the background database of *Arabidopsis thaliana* (UP000006548). Venn diagram was made using InteractiVenn software. Heatmap visualization, and statistical analysis was carried out using the MetaboAnalyst (v6.0) web tool. Two-sided unpaired Student’s t-tests were used to compare the groups statistically. Principal component analysis (PCA) was done using SRplot web tool (Tang et al., 2023). PCA and volcano plots were generated using ggplot2 package in R (v4.3.1). ShinyGO (v0.80) tool was employed to perform over-representation analysis using the KEGG and Gene ontology: Biological process (GO: BP) database for pathway enrichment.

### Oligosaccharide infiltration and DAB staining

Leaf AIR was digested with xyloglucanase (E-XEGP, Megazyme, Ireland) and total sugar was estimated in xyloglucan-rich oligosaccharide fraction by phenol-sulfuric acid method and diluted to 200 µg of total sugar for infiltration in rosette leaves of three-week-old Arabidopsis plants. The infiltrated leaves were subjected to DAB staining in which leaves were dipped in staining solution prepared by dissolving 3,3’-diaminobenzidene tetrahydrochloride hydrate (DAB.4HCl.xH_2_O) (17076, SRL, India) in water and pH was maintained by 200mM disodium hydrogen phosphate (TC507M, Himedia, India) and incubated at room temperature for 12 h. The leaves were transferred to destaining solution [ethanol(3): acetic acid(1): glycerol(1)], and heated at 95°C for 15 min. The pictures were taken and analyzed in ImageJ.

### Pseudomonas syringae infection assay

For the infection, a tipfull of dividing cells of *Pseudomonas syringae pv. tomato* DC3000 was resuspended in 10mM MgCl_2_ and infiltrated in rosette leaves of three-week-old Arabidopsis plants at a density of 1×10^6^ CFU/ml. Leaf discs were collected at day-0 and day-3 post infiltration from the infected leaf area and macerated in 10mM MgCl_2_, which was further serially diluted to 10^-3^, 10^-4^, and 10^-5^ and plated on Pseudomonas agar plates with 100 µg rifampicin and cycloheximide antibiotics. Colonies on plates were counted, and bacterial growth was determined.

### Ralstonia solanacearum infection assay

For preparing the bacterial inoculum, the freshly grown *R. solanacearum* F1C1 strain (Kumar et al., 2016) was grown in BG broth. The secondary culture of F1C1 was then pelleted down at 3155g for 10 min, washed with sterile distilled H_2_O, followed by resuspension in an equal volume of sterile distilled H_2_O such that the bacterial concentration remains 1 x 10^8^ CFU/ml. The culture of *R. solanacearum* strain F1C1 was drench inoculated into the pots having three-week-old Arabidopsis plants. Plants were incubated in an infection chamber pre-set at 28°C, 12/12 h photoperiod, and 80% humidity. The wilting symptoms were recorded, and plants were photographed using a DSLR camera.

## ACKNOWLEDGMENTS

We would like to RCB-ATPC sequencing platform, RCB-Microscopy facility, RCB-Central Instrumentation Facility (CIF) for their support.

## AUTHOR CONTRIBUTIONS

LR performed most of the experiments. Oligosaccharide fingerprinting analysis was performed by AV and MH. Docking studies were performed by VK and ST. GJ and KK performed Ralstonia infection assay. LR with the help of NK, and TKM performed and analysed proteomics experiments. LR and PP wrote the manuscript with inputs from the authors. PP conceptualized, designed, and secured funding for the project. All authors read and agree to publish the manuscript.

## Supplementary material

**Supplementary Figure S1**. *AtGELP53* gene structure with its T-DNA insertion site.

**Supplementary Figure S2**. Enzymatic oligosaccharide profiling of *atgelp53* homozygous mutant.

**Supplementary Figure S3**. Acetyl content analysis in MS grown seedlings.

**Supplementary Figure S4**. Flow chart of sequential extraction of cell wall polysaccharides from alcohol insoluble residue (AIR).

**Supplementary Figure S5**. Structural analysis of *At*GELP53.

**Supplementary Figure S6.** Cell wall sugar composition of *35S::AtGELP53* OE lines.

**Supplementary Figure S7.** Comparison of proteomics studies between wild-type and *35S::AtGELP53* OE line.

**Supplementary Figure S8.** Proteomics data of leaves infiltrated with xyloglucanase-digested oligosaccharides derived from wild-type and *35S::AtGELP53* OE line after 30 min post infiltration by mass spectrometry.

**Supplementary Table S1**. Top seven best structural homologs of *At*GELP53 identified by a DALI search against PDB.

**Supplementary Table S2**. Monosugar composition analysis (mol%) in leaf and stem tissue of wild-type and *At*GELP53 overexpression lines analyzed by ion-chromatography (IC).

**Supplementary Table S3**. Primer list.

