## Supplementary figures and images for "Arabidopsis *At*GELP53 modulates polysaccharide acetylation and defense response through oligosaccharide-mediated signaling"

### Supplemental Figure 1

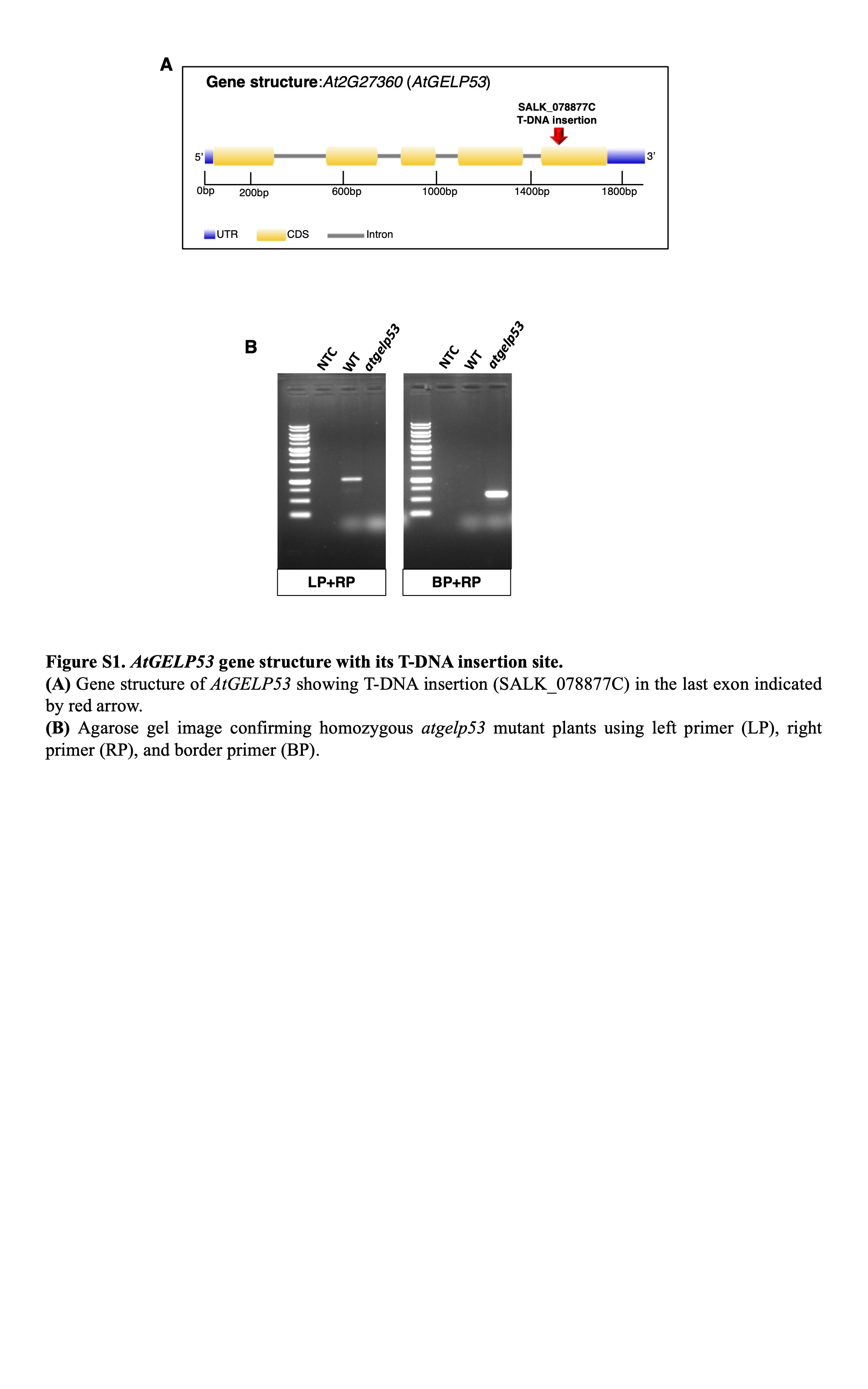

### Supplemental Figure 2

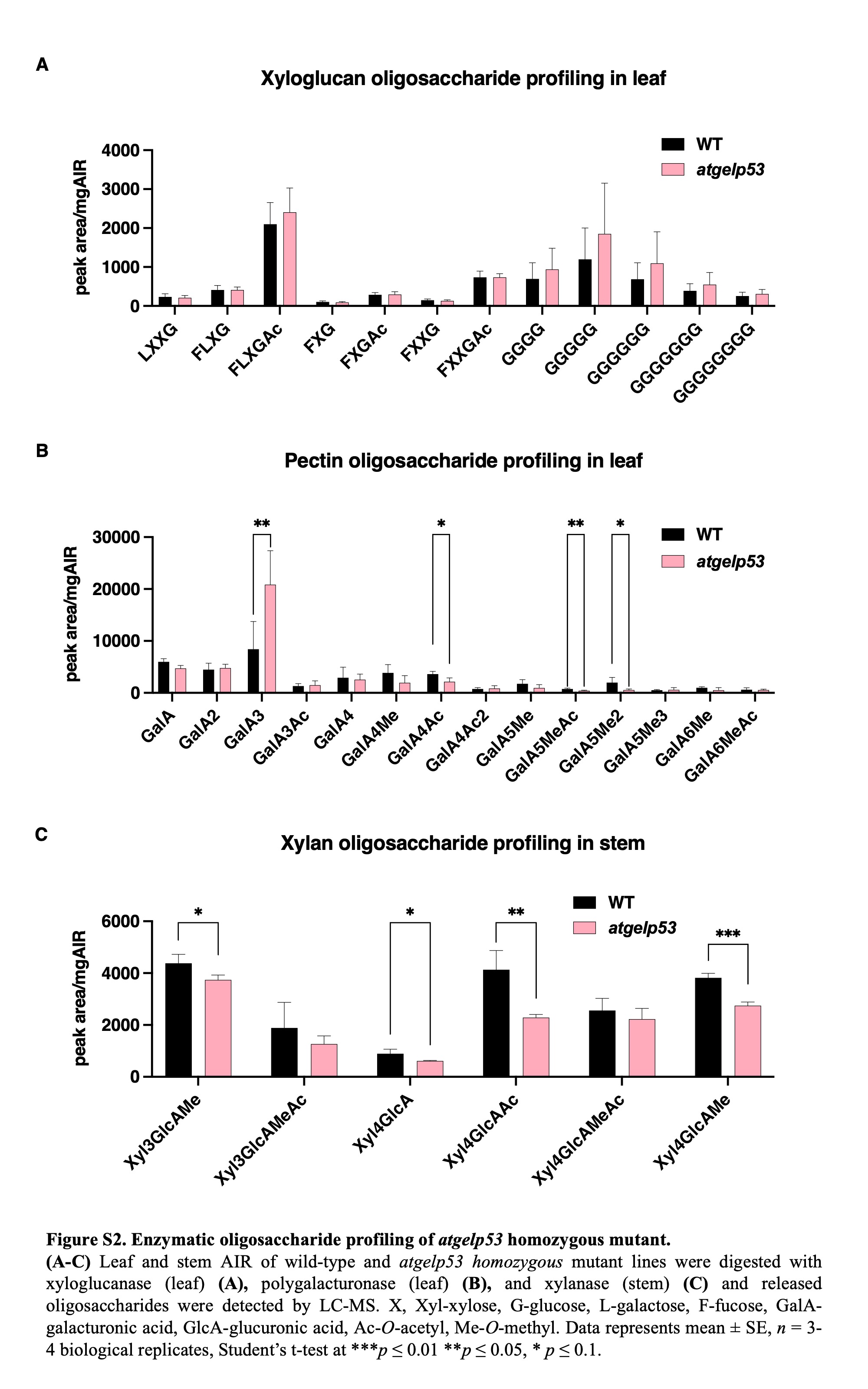

### Supplemental Figure 3

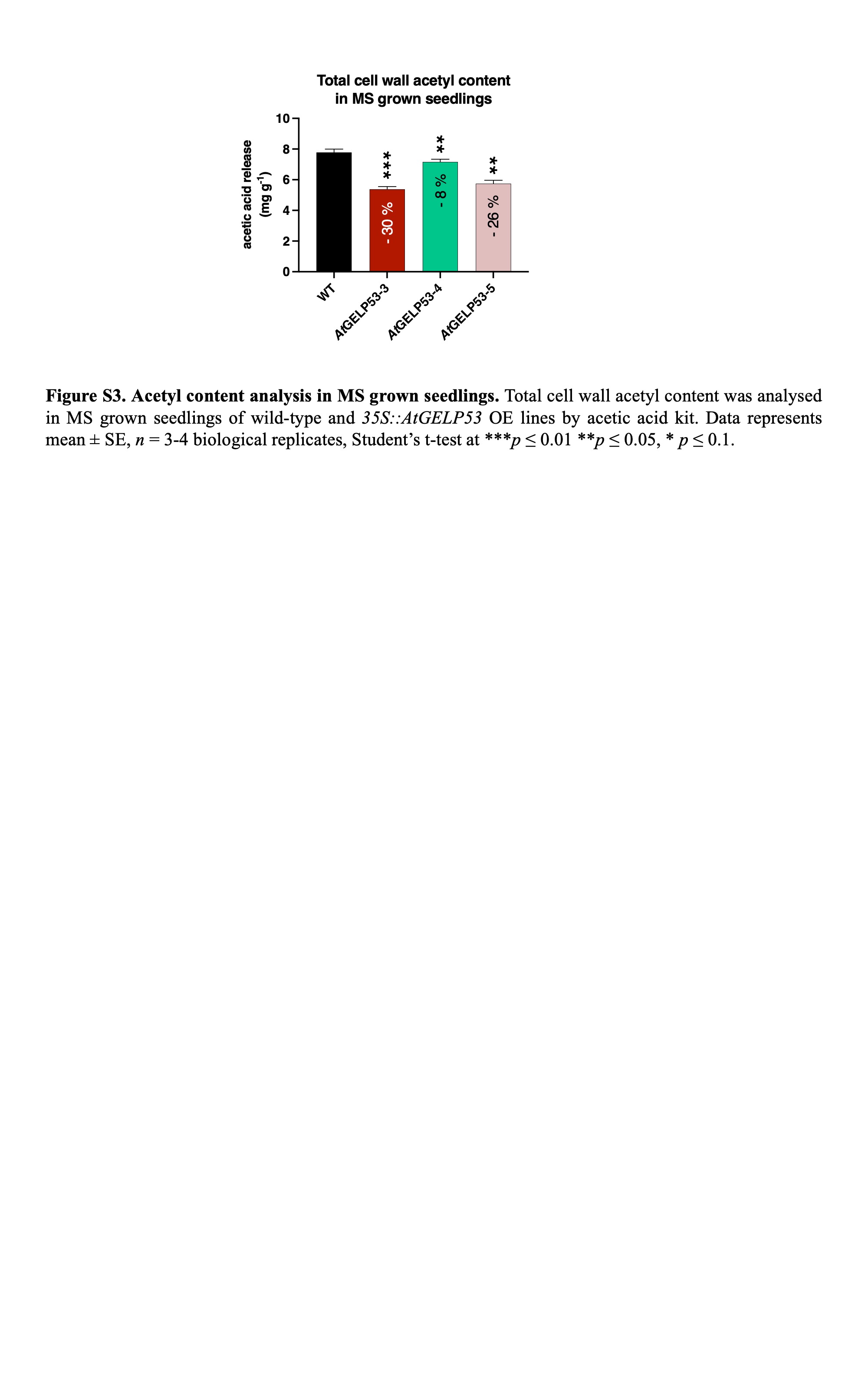

### Supplemental Figure 4

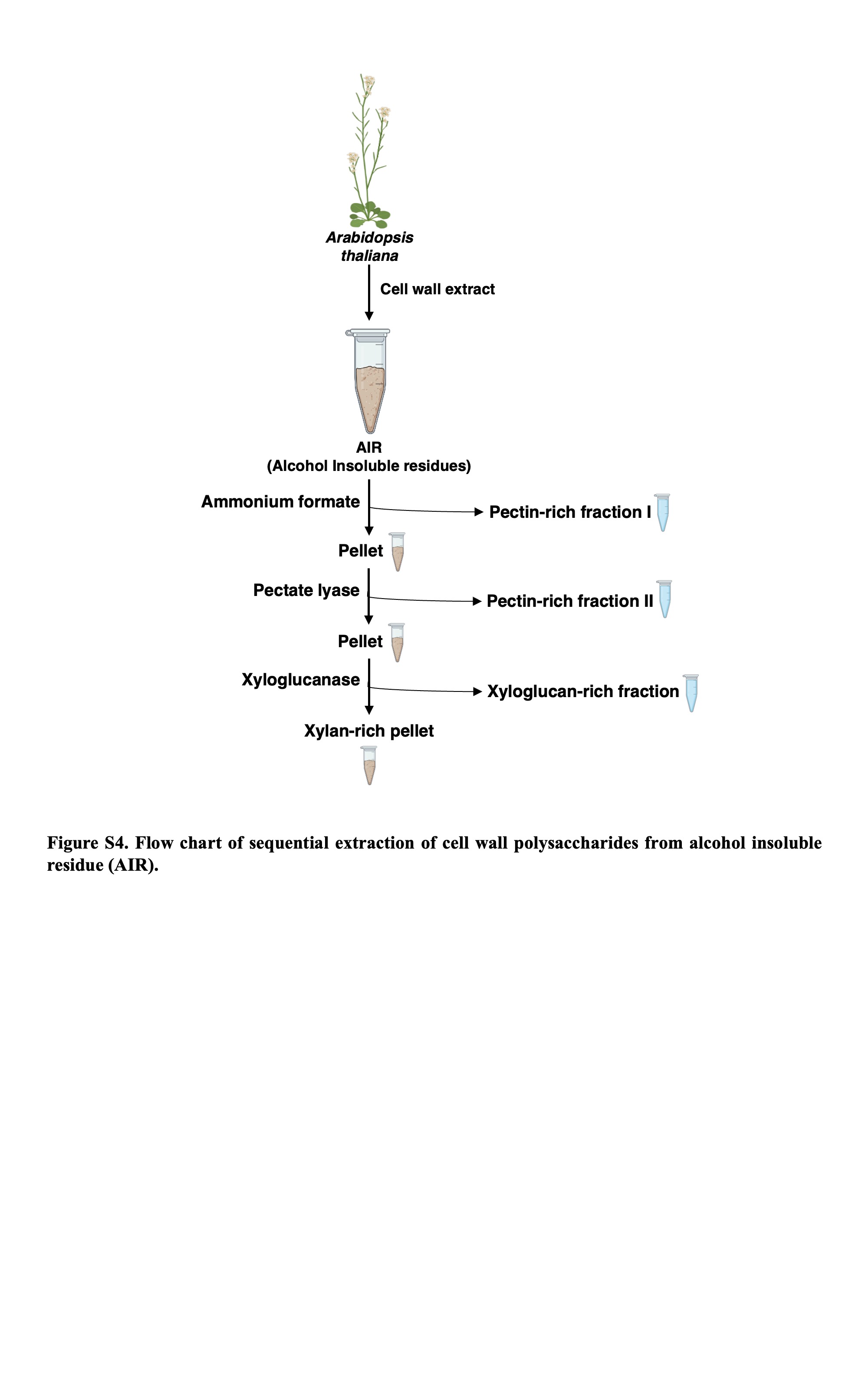

### Supplemental Figure 5

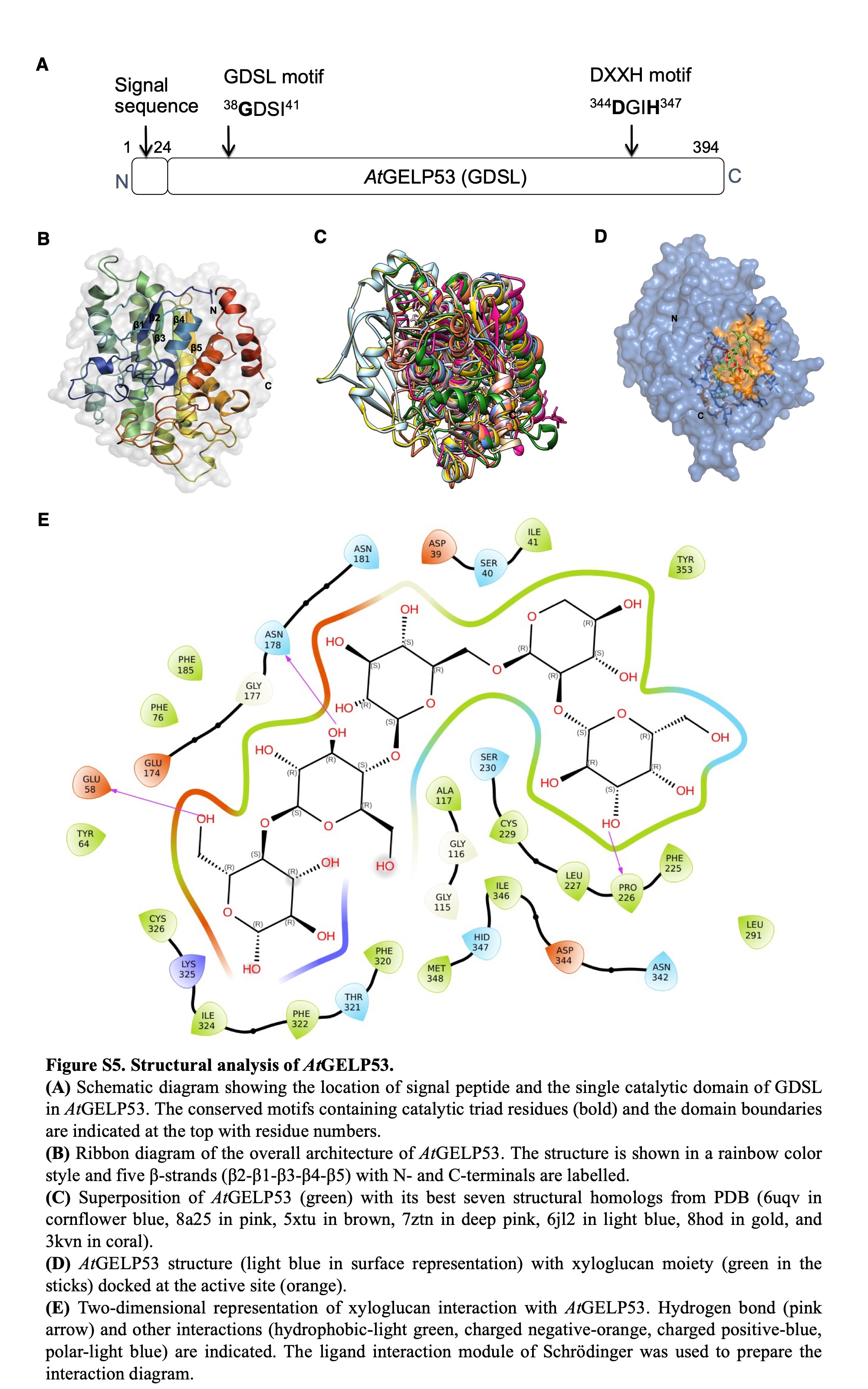

### Supplemental Figure 6

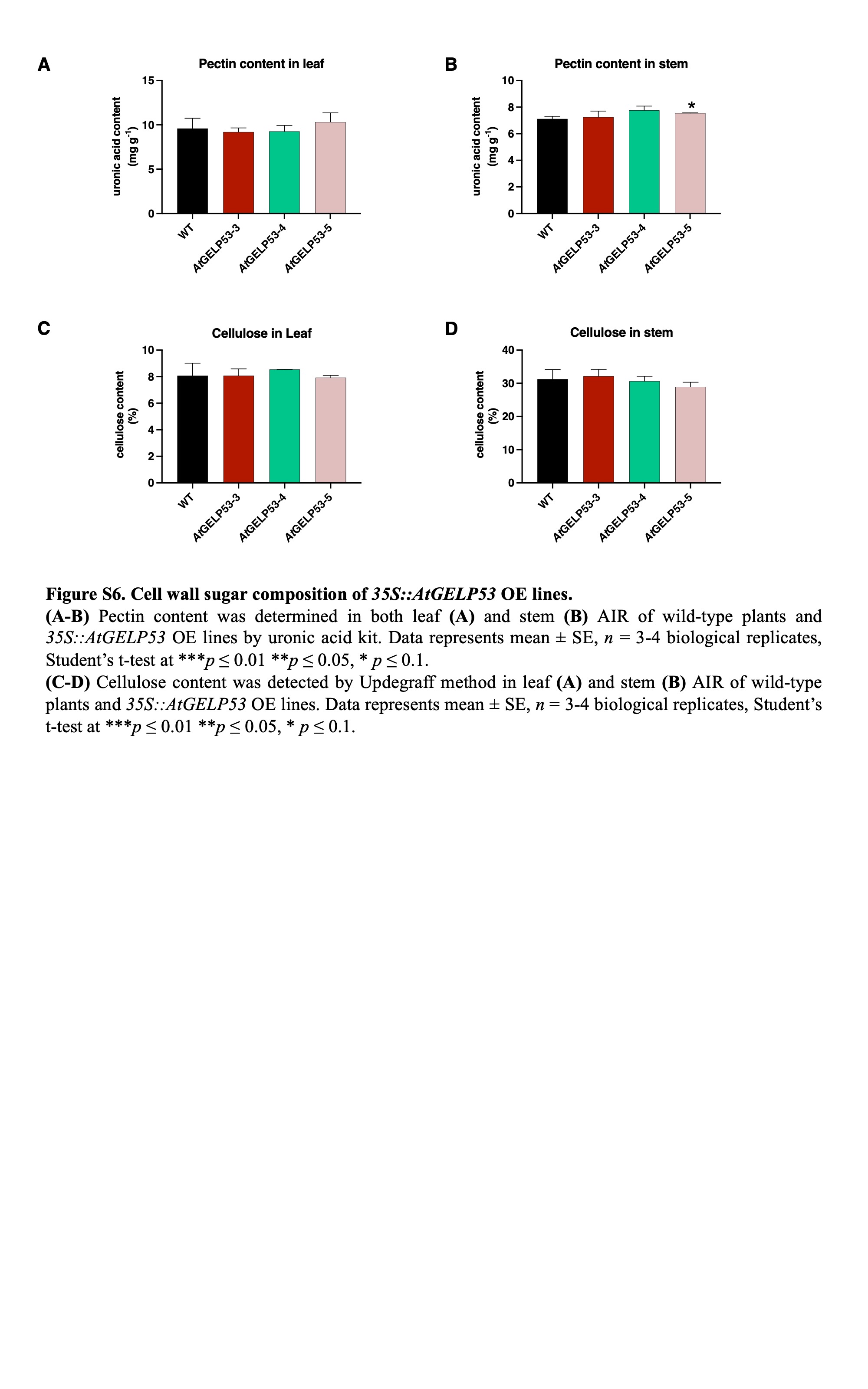

### Supplemental Figure 7

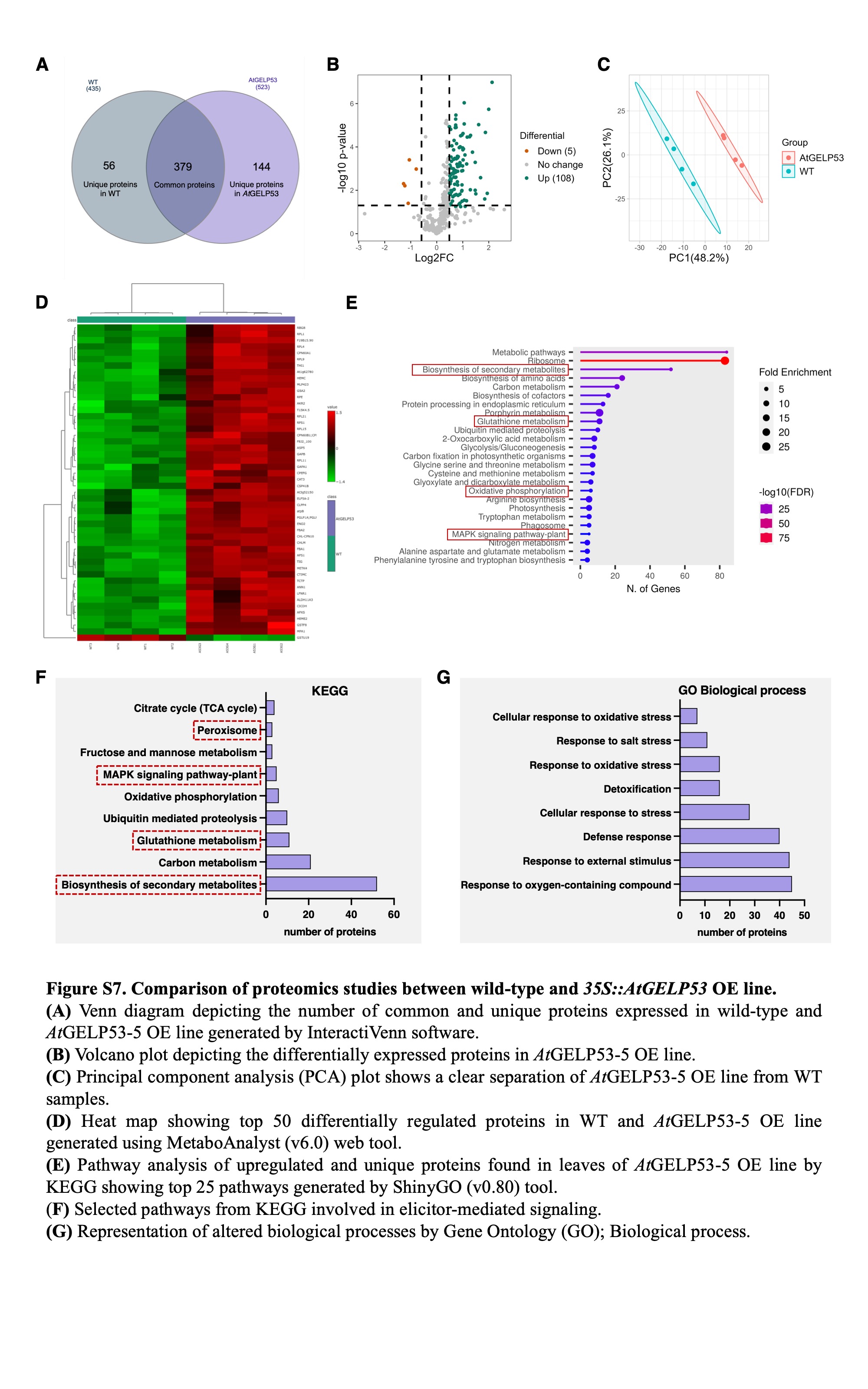

### Supplemental Figure 8

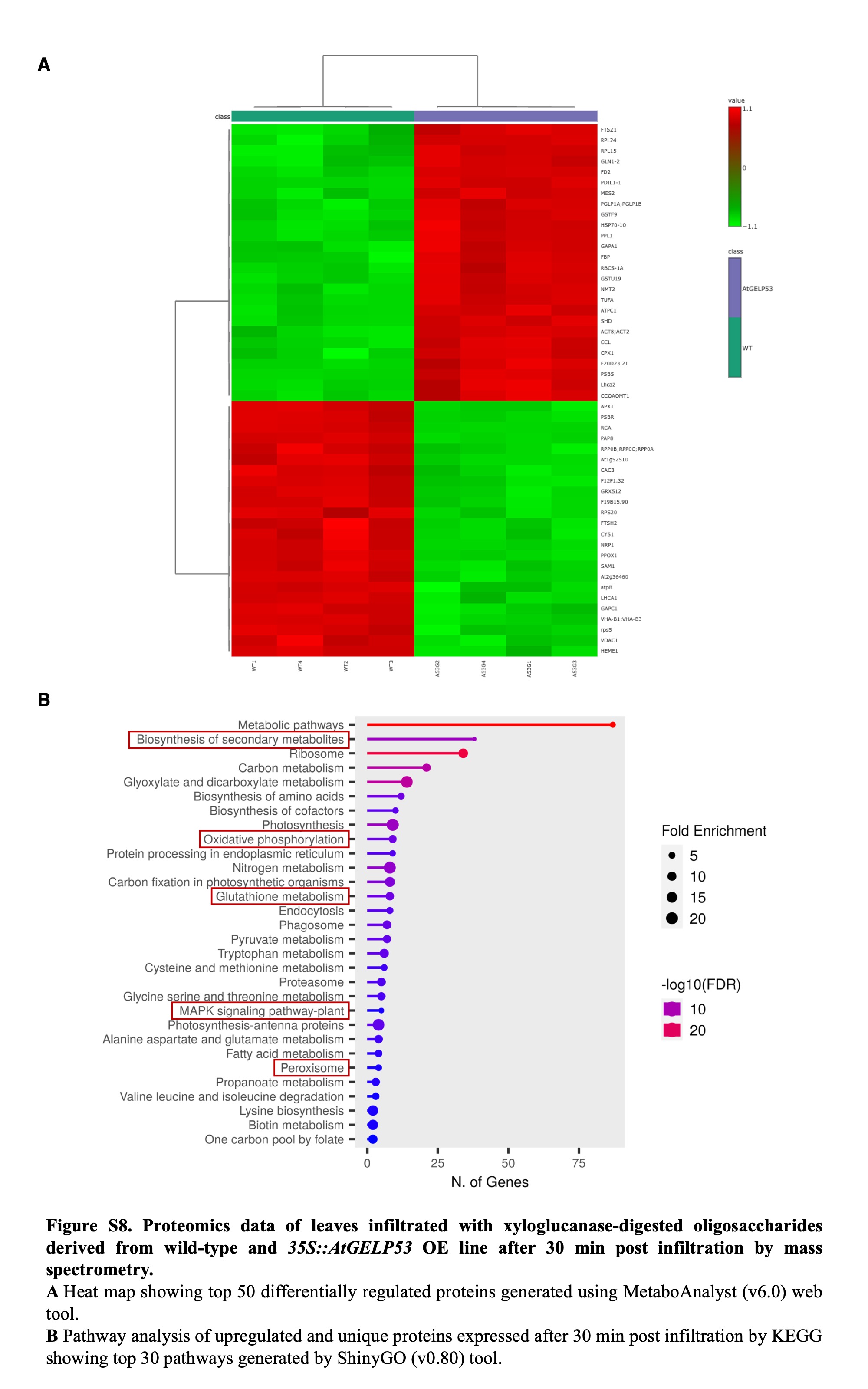
