## Supplemental Tables for "Arabidopsis *At*GELP53 modulates polysaccharide acetylation and defense response through oligosaccharide-mediated signaling"

**Table S1. Top seven best structural homologs of *At*GELP53 identified by a DALI search against PDB.**

| PDB ID | Description | source | Z-score | RMSD  (Å) | No. of aligned residues | Sequence identity (%) |  |
| --- | --- | --- | --- | --- | --- | --- | --- |
| 6UQV | ChoE  acetylcholinesterase | *Pseudomonas aeruginosa* | 30.8 | 2.2 | 269 | 22 | Pham VD et al., 2020 |
| 8A25 | PlaA Lysophospholipase | *Legionella pneumophila* | 29.5 | 2.4 | 269 | 25 | Hiller M et al., 2024 |
| 5XTU | GDSL-esterase | *Photobacterium sp.* J15 | 28.0 | 2.6 | 268 | 23 | Mazlan SNHS et al., 2018 |
| 7ZTN | CE16 acetyl xylan esterase | *Thermothelomyces thermophilus* | 27.1 | 3.1 | 271 | 17 | Pentari C et al., 2024 |
| 6JL2 | VvPlpA phospholipase | *Vibrio vulnificus* | 26.1 | 2.9 | 265 | 23 | Wan Y et al., 2019 |
| 8H0D | VDLT vibrio dual lipase/transferase | *Vibrio alginolyticus* | 25.3 | 7.8 | 258 | 22 | Wang C et al., 2023 |
| 3KVN | EstA autotransporter | *Pseudomonas aeruginosa,* | 24.5 | 3.1 | 268 | 21 | van den Berg B, 2010 |

**Table S2. Monosugar composition analysis (mol%) in leaf and stem tissue of wild-type and *At*GELP53 overexpression lines analyzed by ion-chromatography (IC).** Data represents mean ± SE, n = 3-4 biological replicates, Student’s t-test at ***p ≤ 0.01 **p ≤ 0.05, * p ≤ 0.1.

| Leaf | Xylose | Arabinose | Rhamnose | Fucose | Galactose | Mannose | Glucose |
| --- | --- | --- | --- | --- | --- | --- | --- |
| WT | 7.1 ± 0.3 | 10.4 ± 0.3 | 5.3 ± 0.6 | 2.2 ± 0.1 | 10.2 ± 0.4 | 1.7 ± 0.08 | 62.9 ± 1.7 |
| *At*GELP53-3 | 7.7 ± 0.2 | 10.8 ± 0.3 | 5.2 ± 0.2 | 2.4 ± 0.01  ** | 10.6 ± 0.04 | 2.03 ± 0.1 ** | 61.0 ± 0.1 |
| *At*GELP53-4 | 8.0 ± 0.6 | 11.2 ± 0.5 | 5.2 ± 0.2 | 2.5 ± 0.1  ** | 10.6 ± 0.6 | 1.85 ± 0.06 | 60.3 ± 2.2 |
| *At*GELP53-5 | 7.2 ± 0.7 | 9.8 ± 0.3 | 5.2 ± 0.4 | 2.3 ± 0.1 | 9.7 ± 0.4 | 1.716 ± 0.2 | 63.8 ± 2.3 |

| Stem | Xylose | Arabinose | Rhamnose | Fucose | Galactose | Mannose | Glucose |
| --- | --- | --- | --- | --- | --- | --- | --- |
| WT | 68.3 ± 0.5 | 2.2 ± 0.3 | 3.0 ± 0.1 | 0.7 ± 0.09 | 4.4 ± 0.5 | 5.2 ± 0.03 | 14.7 ± 0.6 |
| *At*GELP53-3 | 66.6 ± 1.02 * | 3.5 ± 0.1  *** | 3.5 ± 0.1  ** | 1.0 ± 0.02 ** | 5.6 ± 0.1  ** | 5.3 ± 0.06 ** | 14.1 ± 1.07 |
| *At*GELP53-4 | 65.6 ± 0.3 *** | 3.2 ± 0.4  ** | 3.4 ± 0.04  ** | 1.0 ± 0.06 ** | 5.5 ± 0.4  * | 5.4 ± 0.06 *** | 15.6 ± 0.7 |
| *At*GELP53-5 | 65.4 ± 1.1 ** | 4.0 ± 0.06 *** | 3.7 ± 0.1  *** | 1.1 ± 0.01 *** | 6.0 ± 0.1  *** | 5.4 ± 0.07 ** | 14.1 ± 1.1 |

**Table S3. Primer list.** F; Forward primer, R; Reverse primer, B; Border primer

| Oligo name | Target gene ID | Name of gene | Sequence (5´-3´) |
| --- | --- | --- | --- |
| PCWL-37 F | AT2G27360 | *AtGELP53* | GGGGACAAGTTTGTACAAAAAAGCAGGCTTCGGATCCATGGCGTCTCAAGATTGTCA |
| PCWL-38 R | AT2G27360 | *AtGELP53* | GGGGACCACTTTGTACAAGAAAGCTGGGTT AAGCTTTCAGCTGTTCATCAAAGAAT |
| PCWL-15 F | SALK_078877C | *AtGELP53* | TTGGATTGGAACGAAAACAAG |
| PCWL-16 R | SALK_078877C | *AtGELP53* | TGAGGAGCCACAAAGATATGG |
| PCWL-29 B | T-DNA | Border primer | ATTTTGCCGATTTCGGAAC |
| PCWL-597 F | AT2G27360 | *AtGELP53* promoter | AAAAAGCAGGCTCAGGAAGAAACAAGGAGGCTAA |
| PCWL-598 R | AT2G27360 | *AtGELP53* promoter | AGAAAGCTGGGTAACGACAGTGATTAAAAAAGT |
| PCWL 127 F | AT3G18780 | Actin2 | TGCTGGATTCTGGTGATGGT |
| PCWL 128 R | AT3G18780 | Actin2 | AATTTCCCGCTCTGCTGTTG |
| PCWL-231 F | AT1G13440 | GAPDH | TTGGTGACAACAGGTCAAGCA |
| PCWL-232 R | AT1G13440 | GAPDH | AAACTTGTCGCTCAATGCAATC |
| PCWL_137 F | AT2G27360 | *AtGELP53* | ATGTTTGACATGGCTGAACG |
| PCWL_138 R | AT2G27360 | *AtGELP53* | CCGAATTTGGATGGTTCTTG |
| PCWL-484 F | AT1G70230 | TBL27 | AATGGAAACCAAACGAATG |
| PCWL-485 R | AT1G70230 | TBL27 | GGGACCAGTAAACCGAGACA |
| PCWL-486 F | AT3G06080 | TBL10 | GTGTTCGAGCCGTCTTTAGG |
| PCWL-487 R | AT3G06080 | TBL10 | TCGAACACATCACACCCACT |
| PCWL-76 F | AT3G55990.1 | TBL29 | GGAACCAATGGGAATCAATG |
| PCWL-77 R | AT3G55990.1 | TBL29 | AGAAAGTCAACGCCTTTCCA |
| PCWL-145 F | AT5G46340 | RWA1 | AATGGAAAGGATGGATGCAG |
| PCWL-146 R | AT5G46340 | RWA1 | GCAAAACGTGCAACAGAGAA |
| PCWL-147 F | AT3G06550 | RWA2 | TCAAACCGAGGAGTGGAAAG |
| PCWL-148 R | AT3G06550 | RWA2 | CCACATCATCTGTGCAAACC |
| PCWL-149 F | AT2G34410 | RWA3 | TCAGCCATGACATCCTTGAA |
| PCWL-150 R | AT2G34410 | RWA3 | CAGACGTAGGCAGCAATGAA |
| PCWL-151 F | AT1G29890 | RWA4 | CAATTACGCCTGGTCAGGTT |
| PCWL-152 R | AT1G29890 | RWA4 | TTTGTTCGTCACCATTTCCA |
| PCWL-842 F | AT3G03210 | AXY9 | CGCGGTTTGTGTTCATCTCA |
| PCWL-843 R | AT3G03210 | AXY9 | TTCCTACGAACAGACTCCGG |
| PCWL-406 F | AT1G10670 | ACLA1 | AAAAAGCAGAGCCCTTGTCA |
| PCWL-407 R | AT1G10670 | ACLA1 | GGCTCGCATTTTAGCAAGTC |
| PCWL-408 F | AT1G60810 | ACLA2 | GTAGCTGGAGGAGGTGCAAG |
| PCWL-409 R | AT1G60810 | ACLA2 | TGAAGGTAGCAGCAACATCG |
| PCWL-410 F | AT1G09430 | ACLA3 | GATGACACTGCTGCCTTCAA |
| PCWL-411 R | AT1G09430 | ACLA3 | AGCTACCATCGTCCAAATGC |
| PCWL-395 F | AT3G53260.1 | PAL2 | TGTCCAACGGTGAGACTGAG |
| PCWL-396 R | AT3G53260.1 | PAL2 | CAAGCTCTTCCCTCACGAAC |
| PCWL-383 F | AT4G36220.1 | F5H | ATGATGGGGATGTTGTCGAT |
| PCWL-384 R | AT4G36220.1 | F5H | ACTCCGTTAAGGCCCACTCT |
| PCWL-391 F | AT1G15950.1 | CCR1 | TATGTGGATGTTCGCGATGT |
| PCWL-392 R | AT1G15950.1 | CCR1 | GGCTTGGCTCTAGGGTTCTT |
| PCWL-776 F | AT2G30490.1 | C4H | ACCGAGCCTGATCTTCACAA |
| PCWL-777 R | AT2G30490.1 | C4H | GGTTGTTTGCTAGCCACCAA |
| PCWL-786 F | AT2G40890.1 | C3’H | TCCCGCTTACCTTACTTGCA |
| PCWL-787 R | AT2G40890.1 | C3’H | TTTTCCATACAGCCGGGTCT |
| PCWL-784 F | AT4G34050 | CCoAOMT1 | TGGCTACGACAACACTCTGT |
| PCWL-785 R | AT4G34050 | CCoAOMT1 | CTGATCCGACGGCAGATAGT |
| PCWL-245 F | AT1G21250.1 | WAK1 | TGCTCGTGCTTTCACAAATC |
| PCWL-246 R | AT1G21250.1 | WAK1 | ATGCAAGAGTTCCAGCGACT |
| PCWL-247 F | AT1G21270.1 | WAK2 | ATTGCCGGACAACTCCATAG |
| PCWL-248 R | AT1G21270.1 | WAK2 | GACGGTGCTCCCATGTAAGT |
| PCWL-249 F | AT1G21240.1 | WAK3 | TCATCCATCGCGATATCAAA |
| PCWL-250 R | AT1G21240.1 | WAK3 | TTCTCGTTCAGAAGCCCTGT |
| PCWL-251 F | AT1G21210.1 | WAK4 | AAGTTGGGACACTTCCGTTG |
| PCWL-252 R | AT1G21210.1 | WAK4 | TGCTCGAAGAATTGTTGTCG |
| PCWL-253 F | AT1G21230.1 | WAK5 | CGCCGCCAAATAGTAAATGT |
| PCWL-254 R | AT1G21230.1 | WAK5 | AACACAGGGAACCTCGTGAC |
| PCWL-792 F | AT1G16130 | WAKL2 | CAATGGCGTCTCATCAGCAA |
| PCWL-793 R | AT1G16130 | WAKL2 | GGAGGAAGTTACGGGAGCTT |
| PCWL-241 F | AT5G54380.1 | THESEUS 1 | GAGGTCGACGAGTCCTCAAG |
| PCWL-242 R | AT5G54380.1 | THESEUS 1 | TTTATAAACGCGGCCAAATC |
| PCWL-243 F | AT3G51550.1 | FERONIA | ACGTTTACACCGGAATCAGC |
| PCWL-244 R | AT3G51550.1 | FERONIA | GCGAGATATCATTCCCTCCA |
| PCWL-436 F | AT2G06850 | XTH4 | TGGCACAGGATTTCAATCAA |
| PCWL-437 R | AT2G06850 | XTH4 | TGTTTCCCTTTCCTCCTGTG |
| PCWL-438 F | AT4G30270 | XTH24 | CAACGGCCAGCTTCTTACTC |
| PCWL-439 R | AT4G30270 | XTH24 | TCATCCCAAGTGGATCCTTC |
| PCWL-806 F | AT4G30280 | XTH18 | CCCAACGAAGCAACCAATGA |
| PCWL-807 R | AT4G30280 | XTH18 | GGCCGTTTGAATCGAGTTGT |
| PCWL-470 F | AT2G14610 | PR1 | TTCTTCCCTCGAAAGCTCAA |
| PCWL-471 R | AT2G14610 | PR1 | AAGGCCCACCAGAGTGTATG |
| PCWL-466 F | AT2G19190 | FRK1 | ACGGGCATAGTTCCACAAAG |
| PCWL-467 R | AT2G19190 | FRK1 | CGTCAAAAGAACGACGATGA |
| PCWL-472 F | AT3G26830 | PAD3 | GGCTGAAGCGGTCATAAGAG |
| PCWL-473 R | AT3G26830 | PAD3 | TCCAGGCTTAAGATGCTCGT |
| PCWL-474 F | AT2G38470 | WRKY33 | GAAACAAATGGTGGGAATGG |
| PCWL-475 R | AT2G38470 | WRKY33 | TGTCGTGTGATGCTCTCTCC |
